# Human *rDNA* Copy Number Is Unstable in Metastatic Breast Cancers

**DOI:** 10.1101/623595

**Authors:** Virginia Valori, Katalin Tus, Christina Laukaitis, David T. Harris, Lauren LeBeau, Keith A. Maggert

## Abstract

Epigenetic silencing, including the formation of heterochromatin, silent chromosome territories, and repressed gene promoters, acts to stabilize patterns of gene regulation and the physical structure of the genome. Reduction of epigenetic silencing can result in genome rearrangements, particularly at intrinsically unstable regions of the genome such as transposons, satellite repeats, and repetitive gene clusters including the rRNA gene clusters (*rDNA*). It is thus expected that mutational or environmental conditions that compromise heterochromatin function might cause genome instability, and diseases associated with decreased epigenetic stability might exhibit genome changes as part of their etiology. We find support of this hypothesis in invasive ductal breast carcinoma, in which reduced epigenetic silencing has been previously described, by using a facile method to quantify *rDNA* copy number in biopsied breast tumors and pair-matched healthy tissue. We found that *rDNA* and satellite DNA sequences had significant copy number variation – both losses and gains of copies – compared to healthy tissue, arguing that these genome rearrangements are common in developing breast cancer. Thus, any proposed etiology onset or progression of breast cancer should consider alterations to the epigenome, but must also accommodate concomitant changes to genome sequence at heterochromatic loci.

**Authors’ Statement:** One of the common hallmarks of cancer is genome instability, including hypermutation and changes to chromosome structure. Using tumor tissues obtained from women with invasive ductal carcinoma, we find that a sensitive area of the genome – the ribosomal DNA gene repeat cluster – shows hypervariability in copy number. The patterns we observe as not consistent with an adaptive loss leading to increased tumor growth, but rather we conclude that copy number variation at repeat DNA is a general consequence of reduced heterochromatin function in cancer progression.

## Introduction

The human genome contains significant amounts of repetitive sequences. Many of the most repetitious – the alphoid satellite repeats, Satellites -I, -II, and -III, and the telomeres – may consist of kilobases up to megabases of nearly-identical repeats that, if damaged, may repair using sister chromatids, homologous chromosomes, non-homologues, or even repeats *in cis* as repair templates. Such events may generate interchromatid crossovers, translocations, acentric/ dicentric chromosomes, repeat expansions and contractions, and/or extrachromosomal circles. Normally, these genome-damaging repair events are disfavored by the packaging of repetitive sequences as constitutive heterochromatin, which potentiates pairing, regulates repair, and inhibits recombination. For example, in *Drosophila* special repair processes disfavor non-allelic crossovers during repair of double-strand breaks in heterochromatin (Chiolo et al. 2011; Ryu et al. 2016). Mutations in *Drosophila* that compromise heterochromatin formation allows recombination within constitutive centric heterochromatin and at telomeres (Weiler and Wakimoto 1995; Perrini et al. 2004), deregulates telomere length (Savitsky et al. 2002; Garcia-Cao et al. 2004), destabilizes repeat gene clusters (Peng and Karpen 2009), and derepresses expression of genes transposed into heterochromatic repeats (Elgin and Reuter 2013). Whether such processes exist in other organisms is not yet know, but mutations in the DNA methyltransferases of mouse and humans compromise heterochromatin formation, leading to hypervariability – predominantly loss – of satellite copy number, and derepression of transposable elements (Damelin and Bestor 2007).

In *Drosophila*, many mutational, developmental, and environmental factors affect heterochromatin stability (Spofford 1976). Reduction through any of these means may lead to instability of the repeat sequences replete in the *Drosophila* genome (Aldrich and Maggert 2015). Some expressed genes are organized as tandem repeats and, for unknown reasons, are generally subject to epigenetic regulation such that the arrays consist of interspersed expressed and non-expressed copies despite identical sequence (Miller and Beatty 1969; McStay and Grummt 2008; Guetg et al. 2010). This is perhaps a strategy to maximize both expression and stability, but it renders such genes particularly sensitive to loss and gain in conditions that reduce heterochromatin formation or function as these gene arrays are only partially packaged as silenced heterochromatin. We and others have observed that the *18S*/*5.8S*/*28S* ribosomal RNA gene cluster (henceforth referred to as the “*45S rDNA*,” from the human nomenclature) is a sensitive locus of copy number changes in *Drosophila* induced by mutation or by ecological conditions (Aldrich and Maggert 2014; Aldrich and Maggert 2015; Gibbons et al. 2015). This is expected to be broadly pleiotropic to a cell since the *rDNA* not only controls rRNA production and translational capacity, but the *rDNA* also mediates other processes regulated by the nucleolus (Kobayashi 2008; Xu et al. 2017; Bughio and Maggert 2019; Salim and Gerton 2019). These processes are not fully investigated, nor is the roster of roles in regulating cell-biological responses mediated by the nucleolus fully enumerated. Recently, hints at function in radiation sensitivity and DNA repair, stress response, metabolic rate, and developmental decisions, have become more concrete. Even vaguer notions of roles, such as in “stability” or “heterochromatin formation” have been confirmed and expanded, suggesting that the pleiotropy of *rDNA* copy number may expand well-beyond the expected impacts on protein synthesis and include many more aspects of nuclear function such as telomere maintenance, transposable element silencing, satellite DNA stability, and others (Paredes and Maggert 2009b; Larson et al. 2012; Zhou et al. 2012).

Heterochromatin formation and regulation is not well understood in humans, especially during disease onset and progression, although there is evidence that some diseases may have heterochromatin loss and repeat instability as part of their complex etiology (Atkin and Brito-Babapulle 1981; Zhang and Adams 2007; Dialynas et al. 2008; Pezer and Ugarkovic 2008; Slee et al. 2012; Ci et al. 2018). Breast cancers, along with leukemias and lymphomas, colorectal cancers, and others, are known to involve expansive genome instability, including chromosome aneuploidies and rearrangements (Richard et al. 2000; Chin et al. 2004; Storchova and Pellman 2004; de Vargas Wolfgramm et al. 2013). These cancers, and others, are expected to show defects in *rDNA* regulation because of the cytological presence of Argyrophilic Nucleolar Organizers (AgNORs), silver-stained multiple or aberrant nucleoli. It is not known whether AgNORs are the manifestation of extrachromosomal circles resulting from damage and incorrect repair of the *rDNA*, as they appear to be in *Drosophila*. If AgNORs, *rDNA* loss, and nucleolar defects share common features, then one may reasonably expect that AgNORs and disease-related changes to *rDNA* copy number may portend changes in stress response, differentiation program, DNA damage response, metabolism, chromosome structural stability, epigenetic instabilities, or other nucleolus-related processes.

In breast cancer, *de novo* mutations and copy number variations are known to exist but have been difficult to quantify or monitor because of the heterogeneity of typical tumors *in situ*, thus much of the mutation and copy number mutational analyses have been investigated using cell cultures which can be made clonal and grown to large numbers (Xu et al. 2017). Studies investigating copy number changes in cancer have so-far analyzed such changes in the context of adaptive advantage by the cancer phenotype (Stults et al. 2009; Wang and Lemos 2017), but find them to be small and variable in scope, and without any phenotypic consequence. Some cell lines may show distinct interline differences in *rDNA* copy number, but it is not known whether these existed prior to, or as a consequence of, culturing (Schawalder et al. 2003; Killen et al. 2009). The large variation in natural and presumably-healthy human *rDNA* copy number (Long and Dawid 1980) also raises the possibility that the “aberrant” copy numbers in these cell lines are merely captured isolates from within natural human population variation (Killen et al. 2009). Although copy number variation analysis is progressing with the advent of new low-copy-number sequencing technologies (Xu et al. 2017), repetitious regions of the genome remain difficult to analyze. Repeated DNAs including the *rDNA* continue to be under-reported in databases or under-investigated in the literature, either because they do not have a tradition of being considered as mutagenic or capable or regulatory function, or because they are refractory to sequencing technologies and assemblies. It is routine to cull these sequences from databases prior to curation or analysis, and even those few reported analyses have not been confirmed as accurate using other methods (Gibbons et al. 2015).

The role of *rDNA* copy number variation in populations or single cells, or any changes in somatic tissues linked to disease risk, onset, or severity, will remain hypothetical until we have an easy way to ascertain *rDNA* copy number from small samples. It is with this intention that these studies were undertaken.

## Results and Discussion

Real-time (quantitative) PCR (qPCR) has been used effectively for *rDNA* copy number determination in *Drosophila* (Paredes and Maggert 2009a; Aldrich and Maggert 2014). The benefit of this approach is that it provides robust and sensitive copy number determination with small amounts of genomic DNA extracted from fresh or fixed tissue, and at low cost. The preparation of tissue is technically simple and rapidly done. We applied this general approach to human tissues, where the sensitivity and low cost allowed investigation of *rDNA* copy number changes in small samples, and from many individuals across large populations.

### Design and Validation of Real-Time Quantitative PCR Primers for Copy Number Determination

Based on validated primer design for quantifying *rDNA* copy number in *Drosophila* (Aldrich and Maggert 2014), we designed six primers to amplify three regions of the human *45S* pre-rRNA transcription unit, two sets directed at sequences in the *18S* region (h18S.1 paired with h18S.2, and h18S.3 paired with h18S.4), and one set directed at the *28S* region (h28S.1 and h28S.2) (Figure 1A). Primer sets amplified approximately 100 base pair regions of the rRNA genes with comparable annealing temperatures (Table I). Primer sets were verified by amplifying *rDNA* segments from whole genomic DNA isolated from blood drawn from an apparently-healthy 46-year-old white male with European ancestry and no known family history of cancer. This has been the lab standard for normalization between samples, experiments, or technical replicates.

**Table I.**
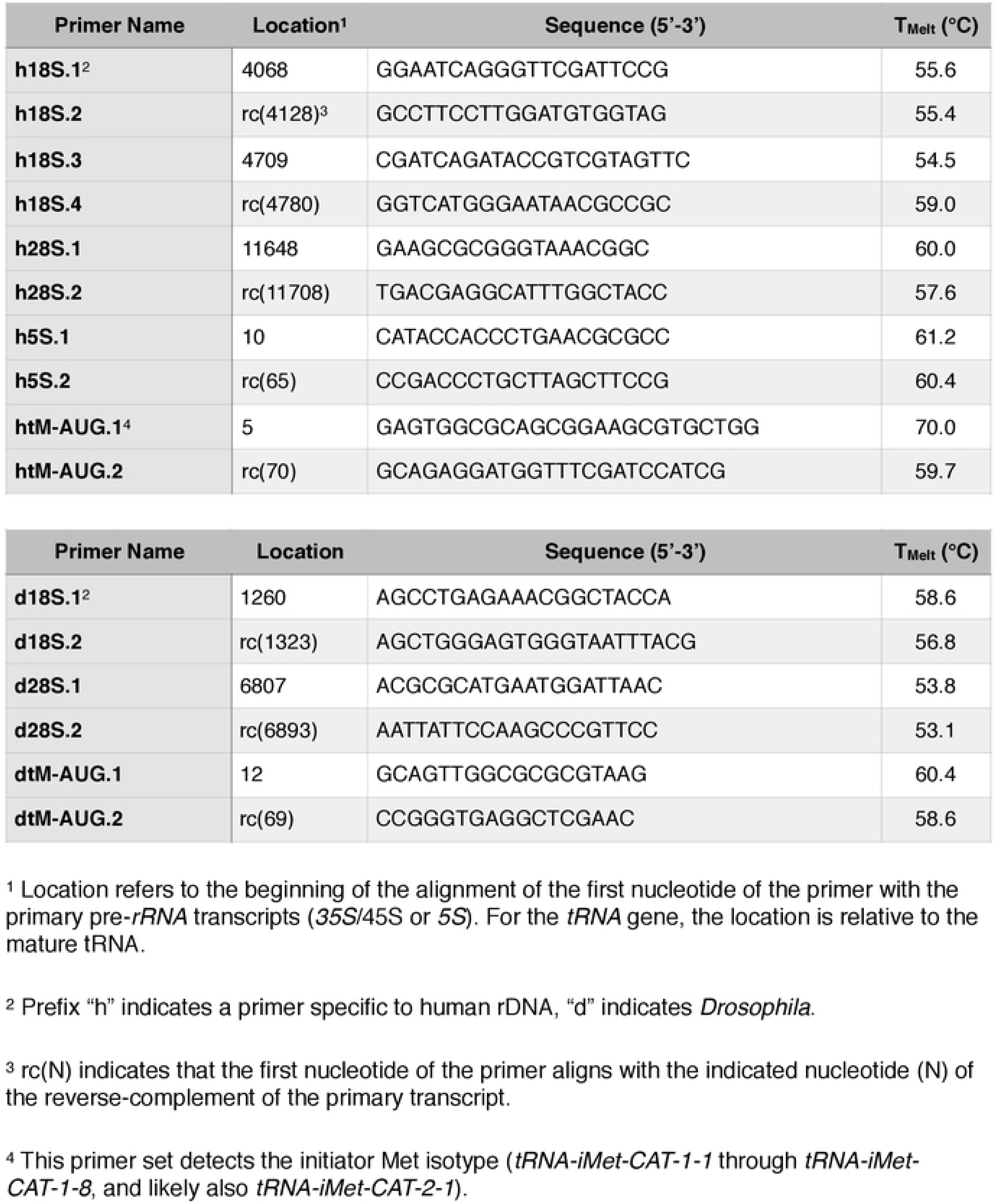
Characteristics of real-time PCR primers in this study

**Figure 1.**
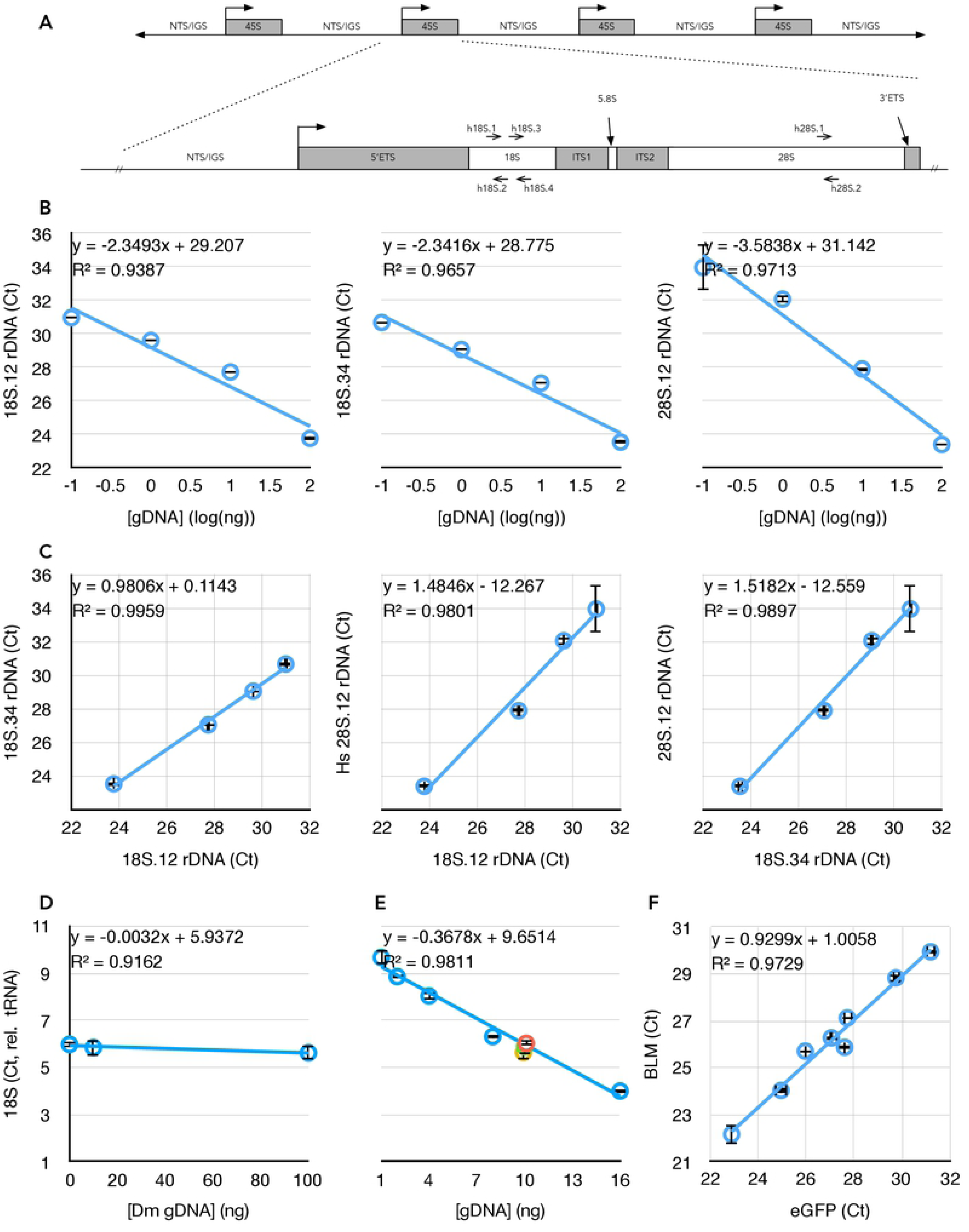
Schematic map of the ribosomal DNA (*rDNA*) Repeat and validation of the Real-Time/quantitative PCR (qPCR) approach taken in this study. **(A)** The *45S rDNA* repeat showing how the structure of the *45S* primary rRNA transcription unit corresponds to the post-processed *18S, 5.8S*, and *28S* rRNA subunits. Locations of the primer sets are indicated. NTS = Non-transcribed Spacer, IGS = Intergenic Spacer, ETS = External transcribed spacer/sequence, ITS = Internal transcribed spacers/sequences. **(B)** Responsiveness of qPCR crossing thresholds (C_t_) to DNA concentration for three different primer sets in (A) (see also Table I), including regression coefficient m (of y = mx+b) and coefficient of determination (R^2^) of the lines. Graphs that do not have numerical ordinal values share an ordinate with other graphs in the same row. **(C)** Correlations between qPCR amplifications of the subunits of the *45S* unit at four different DNA concentrations (from left to right, 0.1 ng, 1 ng, 10 ng, and 100 ng). **(D)** Determination of human *18S rDNA* copy number (using 18S.12 primer set) from 10 ng template DNA, by normalization to human tRNA^Met^ gene copy number (see Table I), in the presence of different concentrations of competing *Drosophila* DNA. **(E)** Determination of human *rDNA* copy number from varied amounts of template DNA, by normalization to human tRNA^Met^ gene copy number, in the presence of competing *Drosophila* genomic DNA such that the total concentration of template DNA was kept constant at 16 ng. Data from (D) are included on this graph as the green, yellow, and orange data points. **(F)** Correlation between copy number determination of enhanced GFP (eGFP) and human Bloom Syndrome helicase (BLM) from cell lines bearing stable integrations of varied copy numbers of a BLM::eGFP fusion transgene. Throughout this figure, error bars indicate standard error of the mean (S.E.M.) for triplicate or quadruplicate technical reactions.

Target *rDNA* sequences are under strong selective pressure and are not known to vary between individuals or between repeat units within individuals (Wellauer and Dawid 1977; Erickson et al. 1981; Wilson et al. 1982; Prokopowich et al. 2003; Richard et al. 2008; Hugerth et al. 2014). Without known exception among eukaryotes, the *18S* and *28S* rRNA subunits of the ribosome are transcribed as a single pre-rRNA transcript from the *45S* rDNA gene, then post-processed in the nucleolus into independent structural ribosomal RNAs (rRNAs). As such, we expected a strong correlation between the two *18S* targets’ and between the *18S* and *28S* targets’ copy numbers, as we are aware of no mechanism or rearrangement in any organism that breaks the fundamental correlation of these two co-transcribed subunit sequences. We validated *rDNA* copy number determination across a 100-fold dilution range for both *18S* and the *28S* target sequences, centered on 10 nanograms per reaction, which was empirically determined to be the most robust in studies of *Drosophila rDNA* copy number (Paredes and Maggert 2009a). Coefficients of determination of 0.94, 0.97, and 0.97, respectively (Figure 1B), and coefficients of determination of 1.0, 0.98, and 0.99 for pairwise comparisons (Figure 1C), confirmed the robustness of the primer sequences in quantifying *rDNA* copy number, even from low DNA concentration samples (1 ng/reaction). These regressions are much stronger than those reported in other studies using other techniques (Gibbons et al. 2015), which is an indicator that qPCR may yield more accurate data than high-throughput sequencing in determining copy number. The ability to perform multiple reactions also allows us to evaluate accuracy in copy number determination. Recalculation of regression excluding the 10 nanogram data points did the least to alter these values, suggesting this concentration is most robust, as in *Drosophila*. Henceforth in this study, unless otherwise indicated, *rDNA* copy number determinations were done with 10 nanograms of genomic DNA per individual reaction, performed at least in triplicate. This amount of template DNA is well within an acceptable range of DNA yield from dried blood cards (average yield > 500 ng/1 cm^2^), fresh unspun blood (average yield > 100 *µ*g/200 *µ*L whole blood), or FFPE sections (average yield ∼ 500 ng/five 10 *µ*m sections). Although we did not test multiple DNA sources from any single individual, the consistency of low- and single-copy number genes (see below) across all samples at all concentrations suggests the source/storage of DNA does not detectably affect copy number determination.

To compare relative *rDNA* copy numbers between different samples, we designed primers to three tRNA genes. Each is multicopy, but their distributed location throughout the genome makes them relatively stable in copy number provided the genomes in question are free from overlapping segmental duplications or deficiencies (*i.e.*, copy number variations), or from chromosomal aneuploidies. To assure that these would be minor sources of error, we also designed primers to multiple single-copy genes amplified in normal cells (Materials and Methods). DNA extracted from the same male peripheral blood and from unrelated primary male human foreskin fibroblast cultures were tested for copy numbers of each tRNA and each single-copy gene.

Specificity of amplified and quantified sequences were confirmed by analysis of the *post-hoc* melt-curves for each of the rRNA and tRNA sequences, each primer set producing a single peak in a graph of the first-order derivative of fluorescence change with respect to temperature. In each case, high-concentration (1.7%) agarose gel electrophoresis was performed and a band of the expected size was the only visible PCR product. Sanger sequencing of five individual reactions further confirmed that the population of amplified DNA was homogenous and corresponded to the desired target sequence. In our hands, the primer sets directed at tRNA^Met^ and 18S.12 performed the best, so unless otherwise indicated these are used for the remainder of the work.

We challenged quantitative amplification of human *rDNA* (and tRNA gene) in two ways. First, we held constant the concentration of human DNA and challenged it with no, 10 nanograms (equivalent mass of DNA), and 100 nanograms (10-fold excess) of *Drosophila* genomic DNA. Given the approximate 10-fold larger genome size of humans than of *Drosophila*, the last condition represents an approximate 100 gene-molar excess of the single copy genes. Although *rDNA* copy numbers vary within both humans and *Drosophila*, they can be considered approximately equal in copy number in broad populations (Long and Dawid 1980). Comparable sequences between these two species are less than perfectly complementary (95% for the 18S. 2 primer, 91% and 64% for the 18S.34 set, and 89% for the 28S.1 primer, Table I), despite being in conserved regions of the rRNAs. Even with molar excess of *Drosophila* genomic *rDNA* target, there was no detectable difference in human *rDNA* quantification (Figure 1D); the coefficient of determination was very high, and the regression coefficient was near-zero (m = −0.003), suggesting that competing *Drosophila* DNA had no measurable impact on quantification of human *rDNA* copy number.

Second, we challenged human *rDNA* amplification by titrating the human DNA to correspond with increased titer of *Drosophila* DNA, keeping the total amount of genomic DNA constant at 16 nanograms. In this configuration, the molar-competition spans from 0 to 150-fold gene-molar excess but at a 10-fold lower DNA concentration than the previous experiment, and again we detected no difference in *rDNA* copy number relative to tRNA normalization (Figure 1E, R^2^ = 0.98). Based on these two competitive experiments, we conclude that the quantification of *rDNA* copy number in humans is remarkably robust to DNA concentration, even when in competition with vast excess of a homologous “contaminating” animal DNA. We expect that contamination by organisms more-diverged than animals (*e.g.*, bacteria, fungi) would be of negligible concern in routine laboratory applications.

A similar analysis was performed using *rDNA* and tRNA gene copy number with three single-copy genes. Without exception, copy numbers of *rDNA* and tRNA relative to these “denominator” copy numbers were consistent across the range of concentrations used above (R^2^ for *Rnmt2* (Gene ID: 1787) was 0.98 over the same concentration range shown in Figure 1B, R^2^ for *Snail2* (Gene ID: 6591) was 0.99, R^2^ for *Bloom Helicase* (Gene ID: 641) was 0.98). Finally, we confirmed the sensitivity and robustness of relative copy number determination of the last gene by using a series of established Bloom Syndrome cell line clones (all derived from a precursor GM08505 strain), each containing stably-transformed (*i.e.*, integrated) *BLM-GFP* fusion genes. Eight sub-lines have different numbers and loci of integrations, but *BLM* and *GFP* sequences always co-varied (R^2^ = 0.97, Figure 1F).

It is critical to emphasize that absolute copy number comparisons of *rDNA*, tRNA gene, and unique genes are not possible. This is the unavoidable expectation, as copy number is derived from qPCR reaction crossing thresholds (C_t_s), which are affected by primer sequence, annealing kinetics and temperature, the length and sequence composition of the inter-primer sequence, subtle sequence biases in SYBR binding, *etc.* For this reason, while comparisons of relative copy number between samples or conditions is valid, absolute determination of copy number of one gene relative to another in any one sample is certainly not. This is clear from the data of Figure 1F, where the Y-intercept of a known 1-to-1 correlation is not zero (b = 1.01).

Relative amplification of single copy and tRNA genes were compared between male blood and genomic DNA extracted from tissue sections from ovaries of three healthy human women. tRNA/single copy genes did not vary, although *rDNA*, telomeric repeats, and satellites did (Figure 2A). This situation is expected given the known variation in these repetitive sequences between individuals in the population. Three genomic DNA preparations from three sequential sections of the same ovarian tissue showed *rDNA* repeat determinations to be robust across technical replicates (Figure 2B).

**Figure 2.**
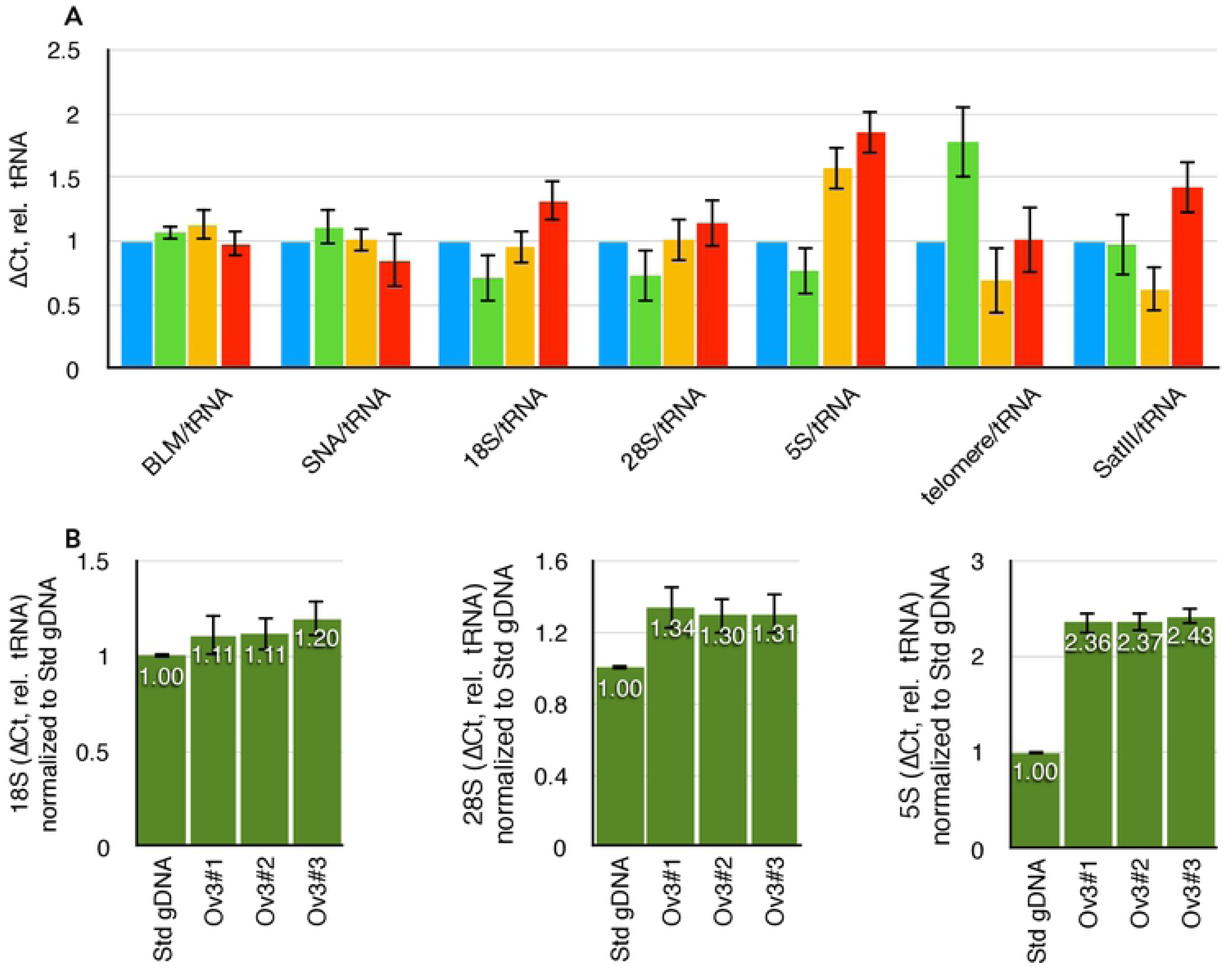
Variation in repeat DNA copy number in different individuals. **(A)** Variation in copy numbers of single-copy genes (*BLM* and *SNA*), three *rDNA* targets (*18S, 28S*, and *5S*), telomeric repeats, and satellite-III, relative to tRNA^Met^. Blue data are from the laboratory standard DNA, and are by definition set to unity. Green, yellow, and red datasets are from three different individual ovaries obtained from the Univeristy of Arizona Tissue Acquisition and Cellular/Molecular Analysis Shared Resource without any identifying information. **(B)** Data obtained from three consecutive sections of one individual ovary. “Std gDNA” is the laboratory standard *human* genomic DNA from peripheral blood. Throughout this figure, error bars indicate standard error of the mean (S.E.M.) for triplicate or quadruplicate technical reactions. S.E.M. from the laboratory standard is pooled into the data from the other individuals’ values.

### Epigenetic Instabilities in Breast Cancer - AgNOR to *rDNA* loss?

Breast cancer is among one of the cancers with clear cytological manifestations of nucleolar instability (Lesty et al. 1992; Bankfalvi et al. 1999; Barwijuk-Machala et al. 2004; Derenzini et al. 2004). The presence of supernumerary argyrophilic nucleolar organizing regions (AgNORs) is interpreted as either fragmentation of nucleoli or the derepression of inactive *rDNA* arrays (or both). In *Drosophila*, presence of supernumerary nucleoli correlates with *rDNA* excision to create extrachromosomal circles of *rDNA* genes (Peng and Karpen 2007), likely a result of derepression, damage, and repair from template copies *in cis* (Guerrero and Maggert 2011; Aldrich and Maggert 2015). Further, the appearance of supernumerary or “fragmented” nucleoli correlates with the severity of *rDNA* loss, as is expected from a simple excision of acentric *rDNA* extrachromosomal circles followed by a cell division (Paredes and Maggert 2009b). Whether AgNORs present in breast cancer tumors correspond to *rDNA* copy number loss in those tumors is not know.

We obtained twenty-nine samples of breast cancer tumors (Table II). Tumors were from invasive ductal carcinomas with evidence for lymph node metastases, and were of various genetic subtypes — Estrogen Receptor positive or negative, Progesterone Receptor positive or negative, HER2 over-expression or not (typed on the 0-3+ scale, where 1-2 were taken as negative), and of varied Ki-67 scores; two were “triple-negative” and had Ki-67 fractions of 90%. In each case, formalin-fixed paraffin-embedded tumor tissues were obtained and two sequential 10 *µ*m slices made. The first was haemotoxilin-eosin stained to define tumor tissue and healthy marginal tissue from the same patient sample. We attempted to normalize areas on the slides with similar cell numbers, avoiding connective tissue and adipose when possible. Tumor and non-tumor areas were marked, and the corresponding areas on the unstained second section were scraped and DNA isolated using filter-binding after xylenes extraction. For each paired sample, we determined *tRNA*^*M-AUG*^ gene, *18S, 28S*, and *5S rDNA* copy numbers. Throughout our samples, when tumor and non-tumor samples are independently analyzed, the *18S* and *28S* remained correlated, further indicating bona fide changes in *45S rDNA* copy number and not artifactual vagaries of DNA extraction or the qPCR reactions.

**Table II.** Demographics of breast samples

| Sample ID<sup>1</sup> | Age<sup>2</sup> | Race<sup>2</sup> | Stage | EST-R<sup>3</sup> | PROG-R<sup>3</sup> | Her2<sup>3</sup> | Ki-67<sup>3</sup> |
| --- | --- | --- | --- | --- | --- | --- | --- |
| 1 | 42 | White | invasive ductal carcinoma + 1 Lymph. metastasis | NEG | NEG | 1+ | 0.9 |
| 2 | 66 | Asian | invasive ductal carcinoma + 1 Lymph. metastasis |  |  |  |  |
| 3 | 79 | White | Multiple foci of invasive ductal carcinoma + 6/19 Lymph. metastasis | POS | NEG | 1+ | 0.02 |
| 4 | 47 | White | invasive ductal carcinoma + 1/23 Lymph. metastasis | POS | POS | 0+ | 0.25 |
| 5 | 57 | White | invasive ductal carcinoma + 2/15 Lymph. metastasis | POS | POS | 0+ | 0.2 |
| 6 | 58 | White | invasive ductal carcinoma + 5/19 Lymph. metastasis | POS | POS | 0+ | 0.3 |
| 7 | 59 | White | invasive ductal carcinoma + 2/10 Lymph. metastasis | POS | POS |  |  |
| 8 | 62 | Other | infiltrative ductal carcinoma + 2/10 Lymph. metastasis | POS | POS | 1+ | 0.1 |
| 9 | 62 | Other | infiltrative ductal carcinoma + 2/10 Lymph. metastasis | POS | POS | 1+ | 0.2 |
| 10 | 56 | White | invasive ductal carcinoma + 1 Lymph. metastasis | POS | POS | 0+ | 0.1 |
| 11 | 85 | White | invasive ductal carcinoma |  |  |  |  |
| 12 | 50 | White | invasive ductal carcinoma + 4/22 Lymph. metastasis | POS | POS | 0+ | 0.15 |
| 13 | 62 | White | invasive ductal and lobular carcinoma + no Lymph. metastasis | POS | NEG | 1+ | 0.7 |
| 14 | 64 | White | invasive ductal carcinoma + 1 Lymph. metastasis | POS | POS | 2+ | 0.5 |
| 15 | 52 | American Indian/Alaska Native | invasive ductal carcinoma + 2/2 Lymph. metastasis |  |  |  |  |
| 16 | 67 | White | invasive ductal carcinoma + 1/13 Lymph. metastasis | POS | POS | 0+ | 0.1 |
| 17 | 87 | White | invasive ductal carcinoma + 1 Lymph. metastasis | NEG | NEG | 3+ | 0.7 |
| 18 | 43 | White | invasive ductal carcinoma + 1 Lymph. metastasis | NEG | NEG | 3+ | 0.4 |
| 19 | 60 | White | invasive ductal carcinoma + 1 Lymph. metastasis | POS | NEG | 0+ | 0.3 |
| 20 | 71 | White | invasive lobular carcinoma + 2/8 Lymph. metastasis | POS | POS | 1+ | 0.25 |
| 21 | 83 | White | invasive ductal carcinoma | POS | NEG | 1+ | 0.5 |
| 22 | 50 | White | invasive ductal carcinoma + 1/14 Lymph. metastasis | POS | NEG | 1+ | 0.15 |
| 23 | 43 | White | invasive ductal carcinoma + 1 Lymph. metastasis | NEG | NEG | 0+ | 0.9 |
| 24 | 57 | White | invasive ductal + no Lymph. metastasis | POS | POS | 2+ | 0.5 |
| 25 | 77 | White | invasive ductal carcinoma + 1 Lymph. metastasis | POS | NEG | 3+ | 0.05 |
| 26 | 50 | White | invasive ductal carcinoma + 5/15 Lymph. metastasis | POS | NEG | 1+ | 0.1 |
| 27 | 51 | Other | invasive ductal carcinoma + 7/12 Lymph. metastasis | POS | NEG | 0+ | 0.5 |
| 28 | 56 | White | invasive lobular carcinoma + 1 Lymph. metastasis | POS | NEG | 0+ | 0.5 |
| 29 | 62 | White | invasive ductal carcinoma + 1 Lymph. metastasis | POS | POS | 1+ | 0.3 |

<sup>1</sup> No identifying information is presented here; Sample ID refers to a serial nomenclature for this study alone, and connection of these numbers to de-identified University of Arizona College of Medicine sample numbers are curated in the laboratory of Keith A. Maggert.
<sup>2</sup> Self-identified.
<sup>3</sup> EST-R = Estrogen Receptor; PROG-R = Progesterone Receptor; Her-2 = Her-2, using the 0-3+ scale, with 0-1 taken as negative and 2-3 taken as positive; Ki67 = fraction of cells positive for Ki-67.

By analyzing *28S*/*tRNA* as a function of *18S*/*tRNA* separately for both tumor (Figure 3A, red) and non-tumor (blue) samples, we could conclude that the expected *28S*-*18S* correlation exists and is of equal slope (regression coefficients of 1.83 for tumor and 1.66 for non-tumor, both statistically indistinguishable from the slope of 1.5 representing linearity, as in Figure 1C) and correlation (R^2^ = 0.74 for tumor and 0.53 for non-tumor) for both types of tissues. The difference in coefficients of determination is likely due to a larger experimental error in the tumor-derived samples. Thus, we conclude that the overall *rDNA* array structure remains unaffected in tumors, suggesting that any changes in copy number would involve addition or subtraction of whole *45S* rRNA genes, rather than amplification/loss of the *18S* or *28S* independently. We also detected that the relative *rDNA* copy numbers span the same ranges, with no strong evidence for any bias in the spread of values in tumor or non-tumor samples (Figure 3B). But it is likely that the high variation in copy numbers between individuals might obscure any differences in *rDNA* copy numbers within individuals as a result of the disease.

**Figure 3.**
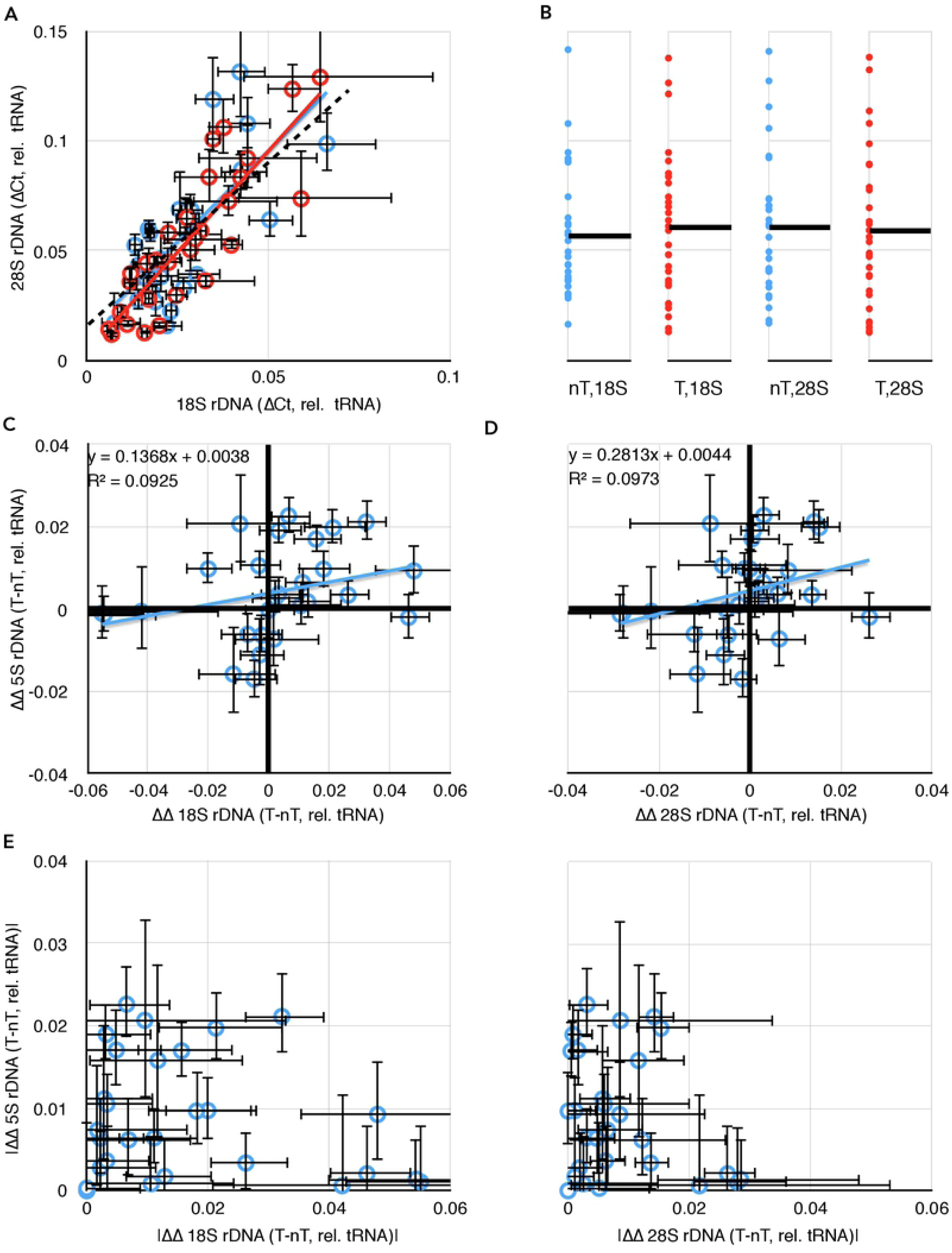
Comparisons of *rDNA* copy number changes in breast cancer tumors. **(A)** Correlation of *18S* and *28S rDNA* copy numbers is retained in both tumor (red) and non-tumor (blue) samples. **(B)** Data from (A) projected into one dimension to visualize the spread of individual data points. Black lines show the means. “nT” = non-tumor, “T” = tumor. The ordinal scale is shared with (A). **(C)** Plot of differences in *18S* and in *5S rDNA* copy numbers between tumor and non-tumor samples from the same individual. Black heavy lines highlight 0,0 origin, indicative of no change between *5S* and/or *18S* copy number between tumor and non-tumor; deviation from the origin is indicative of differences in one or both copy numbers. **(D)** As in (C), but comparing differences in *28S* and *5S rDNA* copy numbers. **(E)** Replotting of the absolute value of the data from 3C and 3D to highlight the discordance in the extent of changes to the *5S* and *45S rDNA* copy numbers. Throughout this figure, error bars indicate standard error of the mean (S.E.M.) for triplicate or quadruplicate technical reactions.

To more sensitively detect changes in *rDNA* copy number, we analyzed the data from Figure 3A as paired (tumor and non-tumor from the same individual biopsy) samples. We considered *5S rDNA* copy number changes by plotting the difference between tumor *5S*/*tRNA* and non-tumor *5S*/*tRNA* (*5S*^T^/*tRNA*^T^ - *5S*^nT^/*tRNA*^nT^) as a function of the differences between *18S rDNA* copy number changes (*18S*^T^/*tRNA*^T^ - *18S*^nT^/*tRNA*^nT^) (Figure 3C) or *28S rDNA* copy number changes (Figure 3D). In both cases we could clearly detect individuals with altered *rDNA* copy numbers (both *18S*/*28S* and *5S*) in tumors compared to non-tumors as those data that deviated from the 0,0 origin. In about half, a clear difference in *rDNA* copy number was detectable between tumor and non-tumor tissue from the same patient. For example, in ten (of 29) samples, the copy number of *28S rDNA* was larger in the tumor relative to the non-tumor, with an average gain of 26% and a population deviation of 15%, and in six (of 29) it was smaller, with an average loss of 22% ± 11%. For the *5S*, ten samples showed an increase in *rDNA* copy number (21% ± 9%), and four showed a decrease (16% ± 9%). As expected from Figure 3A, differences in *18S* and *28S* copy number retained their linear relationships with R^2^ = 0.83 and a regression coefficient of 1.84.

We plotted the absolute value of differences in *5S* as a function of differences in *18S* or *28S* (Figure 3E) to demonstrate that changes in the copy numbers of rRNA genes in these two clusters are not themselves correlated positively or negatively (R^2^ of 0.02 and 0.04, respectively). Thus, it is equally likely in any given tumor sample with an increase in *45S rDNA* copy number to have an increase or a decrease in *5S* copy number, and decreases in *45S* are not enriched for either increases or decreases in *5S*. These results suggest that the changes to the *45S* and *5S rDNA* clusters are independent, and the degrees of changes are uncorrelated. Our finding adds illuminating detail to a previous report (Wang and Lemos 2017), which showed increases in *5S* and decreases in *45S* copy numbers in multiple cancers (including breast cancer) but did not analyze co-relation in the same individual. We detect hyper-variability rather than uniform increases or decreases, indicating that the reported correlation between *45S* and *5S rDNA* copy numbers (Gibbons et al. 2015) is regulated in a way that is ineffective in breast cancer tissues. This possibility would be a striking departure from the expected biology of the *45S* and *5S rDNA* concerted copy number maintenance, and might serve as a powerful diagnostic for breast cancer onset or progression.

That some samples had higher *rDNA* copy numbers, and others had lower, suggests that *rDNA* arrays are subjected to general instability with losses and gains both occurring. This argues against developmental differences, since it seems likely that developmentally-programmed changes to *rDNA* copy number would be uniform in direction, if not in direction and degree. Similarly, it seems unlikely that selective pressures would enrich for tumors with both increases and decreases in copy number if there is a growth (or cancer) advantage to either losses or gains in *rDNA* copy number. Instead, we find these data most easy to reconcile as a result of destabilization of repeat copy number in general. This assertion, and our data, are consistent with those recently published by Xu and colleagues (Xu et al. 2017), and by Wang and Lemos (Wang and Lemos 2017), although both of those studies analyze their data in the context of a cancer adaptive phenotype for *rDNA* copy number changes. These researchers showed that in tumors or samples derived from leukemias and lymphomas, medulloblastomas, osteosarcomas, and esophageal adenocarcinomas, although the average *rDNA* copy number was reduced, there were clear cases of individuals with increased copy numbers.

Those studies reported much larger changes in copy number than did we, but some of the data in those studies indicate copy numbers that were surprisingly low (in some cases, fewer than 5 copies of the *5S* or widely-discordant *18S, 5.8S*, and *28S* copy numbers), indicating their approach may not have generated interpretable absolute copy numbers (see below).

Despite the heterogeneity in genetic subtypes (Table II), we could detect no correlation between *rDNA* variability and any of the genetic markers as categorical variables, or with Ki-67 as a continuous variable (Figure 4). We conclude from this that it is unlikely that these factors, or the known genome instability in triple-negative breast cancers, are driving *rDNA* instability. Rather, we favor the interpretation that the universal loss of heterochromatin function in breast cancers is the cause.

**Figure 4.**
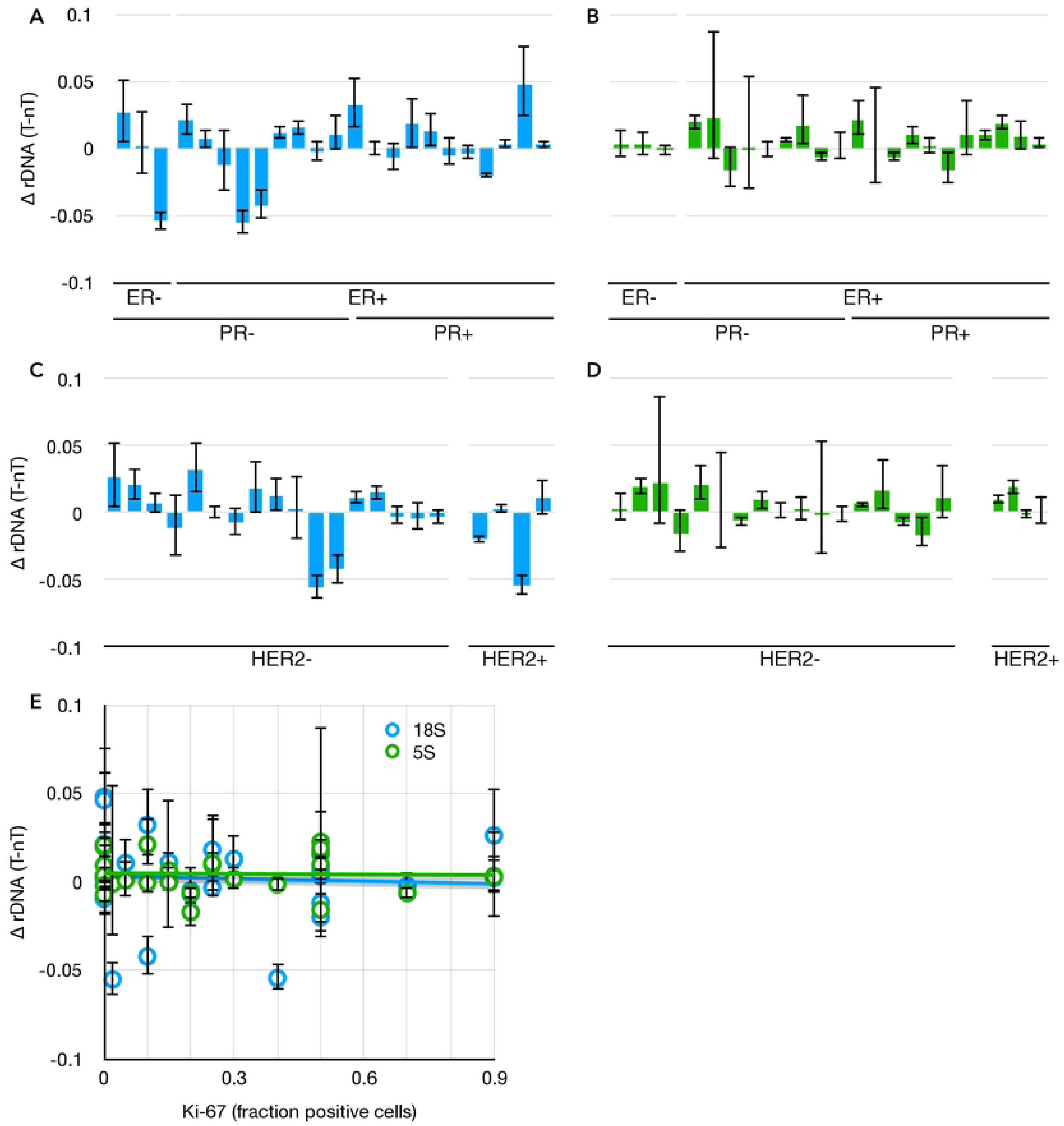
*18S* and *5S rDNA* copy number changes as a function of breast cancer genetic subtype. **(A)** Differences in *18S rDNA* copy numbers between paired tumor and non-tumor samples as a function of categorical grouping of Estrogen Receptor Negative (ER-) and Positive (ER+), and Progesterone Receptor (PR). Data from Figure 3C. **(B)** As in (A), but with *5S rDNA*. **(C)** As in (A), but categorically grouped by *Her2* expression phenotype. Grades 0-1+ were called “negative” and 2-3+ were called “positive.” **(D)** as in (E), but with *5S rDNA*. **(E)** Differences in *18S* (blue) and *5S* (green) *rDNA* copy numbers between paired tumor and non-tumor samples as a function of Ki-67 expression, R^2^ (*18S*) = 0.003, R^2^ (*5S*) = 0.001. Data and S.E.M. are from Figure 3C.

### Variations in Telomeres and Satellite Repeats

The *rDNA* repeats are regulated and stabilized by the formation of heterochromatin (Guetg et al. 2010). We expected that if the underpinning defect in cancer cells is to heterochromatin function, rather than specific regulation of the *rDNA*, then other repeat-sequences would also be affected. To test this, we analyzed “first-order” (repeat-to-tRNA) changes in other repeat DNA sequences: the telomeric repeats and satellite-III sequences. The former are simple and non-transcribed, and altered heterochromatin structures at chromosome ends contributes to telomere length misregulation and allows alternative lengthening of telomeres (Jiang et al. 2009; Jiang et al. 2011) or *de novo* capping by heterologous sequences (Perrini et al. 2004). The latter are stress-responsive transcribed pericentric repeats (Jolly et al. 2004; Valgardsdottir et al. 2005; Jolly and Lakhotia 2006).

We analyzed the telomeric repeats using an approach modified (Aldrich and Maggert 2014) from Richard Cawthon (Cawthon 2002; Cawthon 2009). Telomere repeat copy number in tumor DNA was plotted as a function of copy number in non-Tumor DNA from the same biopsy (Figure 5A). It is clear that deviation from the diagonal (dashed line) is statistically distinguishable from the regression line, but is moderate-to-non-existent in extent (uniformly less than 2%, dotted lines). The coefficient of determination is very low (R^2^ = 0.33), and even these very-small effects are attributable in part to individual variation, rather than the tumor phenotype. Correspondingly, the coefficients of variations (C.V.s) do not differ, C.V. is 0.8% for the non-tumor samples, and 0.6% for the tumor samples. These results are consistent with the relatively small changes in telomere length in breast cancers, despite the common reactivation of telomerase in this cancer type (Bednarek et al. 1997; Nawaz et al. 1997; Umbricht et al. 1999; Rha et al. 2009).

**Figure 5.**
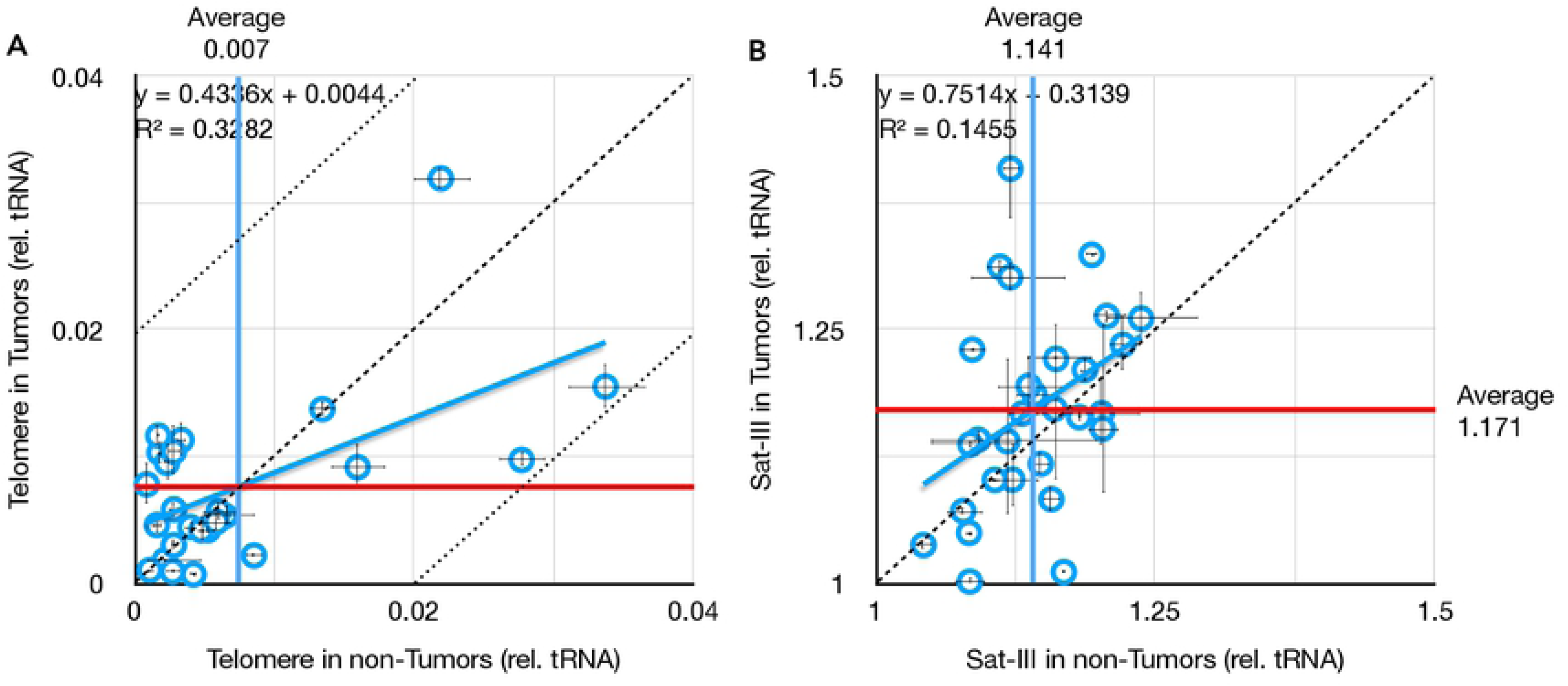
Comparison of telomere and satellite repeats in paired tumors and non-tumor tissues. **(A)** Very weak correlation between telomeric repeat copy number in tumors versus non-tumors from the same individuals. Mean copy numbers are indicated (mean for non-tumors, vertical blue line, is 0.007; mean for tumors, horizontal red line, is 0.008). X- and Y-axis values are percent difference from the lab standard; dashed diagonal is no difference between tumor and non-tumor (y = x), and dotted diagonals demarcate a 2% difference (y = 1x ± 0.02). **(B)** No correlation between Satellite-III in tumors versus non-tumors from the same individuals. Throughout this figure, error bars indicate standard error of the mean (S.E.M.) for triplicate or quadruplicate technical reactions.

For the satellite-III DNA, we used techniques designed for *Drosophila* satellite sequences (Aldrich and Maggert 2014). Three trends were evident upon analysis (Figure 5B). First, tumors exhibit increased variation within the population of 29 individuals analyzed here. Specifically, the coefficient of variation nearly doubles in tumors (C.V. is 4.3% for non-tumors, and 8.3% for tumors). Second, in general, tumors possess more Sat-III copies than do non-tumor tissues from the same individual (range of 1.04 to 1.24 for non-tumors, and range of 1.00 to 1.41 for tumors). Third, the average Sat-III copy numbers were unchanged in tumors compared to non-tumors when taken as populations (mean was 1.14 for non-Tumor samples and 1.17 for tumor samples). These latter two points seem contradictory, highlighting one of the enduring problems of analyzing repeat copy number changes in large populations, namely that copy number is so variable between individuals that even significant changes within individuals are often shrouded when (even small) populations are analyzed (*e.g.*, (Wang and Lemos 2017)). But it is clear from analyzing data from individuals that there are just as many decreases in Sat-III number as increases, and that the increases are larger in scale (+7% ± 7%, range of 1% to 25%) than are the decreases (−4% ± 4%, range of −1% to −13%).

We interpret these data as further support of our interpretation that the proximate cause of genome instability at repeats is a reduction in epigenetic stability of repeat sequences in general. And, as with the *rDNA*, there was no clear trend toward gains or losses of repeat number, merely the occurrence of increased variability. While this may ultimately derive from loss of the *rDNA* itself (Kobayashi 2011), the alternative that a proximate cause of epigenetic instability affecting all repeats lies elsewhere cannot be ruled out. These results indicate that repeat DNA in general, including *rDNA* specifically, are unstable in progressing breast cancer cells.

### Assessing Whether *45S*-*5S* is a Risk Factor for Breast Cancer

Blood is of mesodermal origin, and most of the cells in the breast cancer tumors are of endodermal origin, so these two cell types last shared a common cellular ancestor prior to gastrulation. This allowed us to probe whether we could detect *rDNA* copy number polymorphisms that prefigure later epigenetic instability, as a metric for early risk of later development of breast cancer. This hypothesis derives credibility from the demonstration that naturally- and experimentally-low *rDNA* copy number at fertilization result in adult epigenetic instabilities in model systems, and the finding that loss of *rDNA* leads to cell-autonomous defects in epigenetic stability (Paredes and Maggert 2009b). Blood from the patients from which breast cancer tumors were derived was not available, but the commonality of response (differences between tumor and non-tumor) suggested that if *rDNA* copy number detects prefigured breast cancer in individuals, it might be detected in blood. We therefore obtained genomic DNA from 51 patients involved in a clinical study designed to ascertain breast cancer risk alleles; about half were diagnosed with breast cancer, and none were from the same immediate families. We screened these samples for *rDNA* copy number in the attempt to find if uniformly low, uniformly high, or notably altered *45S*-to-*5S* ratio of *rDNA* copy number was correlated with cancer diagnosis.

The known broad variance in *rDNA* copy number, both *18S* and *28S* (Figure 6A), argued against this simple hypothesis, and against the possibility that *rDNA* copy number could be used as a pre-clinical screen for cancer risk. However, we did discover that the *5S* copy number in all of the study participants was low and did not vary much between patients (Figure 6B, note the much smaller range of the abscissa compared to the ordinate), and showed no detectable correlation with *45S* copy number (R^2^ is 0.08 for patients with a cancer diagnosis, and is 0.03 for non-diagnosed patients). It is of some debate whether *45S* and *5S* copy numbers are equally variable. Analysis of few family lineages has demonstrated that the copy number of *5S* appears more stable than the *45S* (Stults et al. 2008), however the published correlation between *45S* and *5S* copy number in healthy people is strong and robust even between different racial groups worldwide, and under experimental perturbation (Gibbons et al. 2015), suggesting that both copy numbers must be variable. The latter studies reported an approximately 10-fold variance in *45S* and 40-fold variance in *5S rDNA* copy numbers in European men and women (135 of 201 of our samples are non-hispanic white women), while we found only a 1.6-fold variance in the medial 90% of samples.

**Figure 6.**
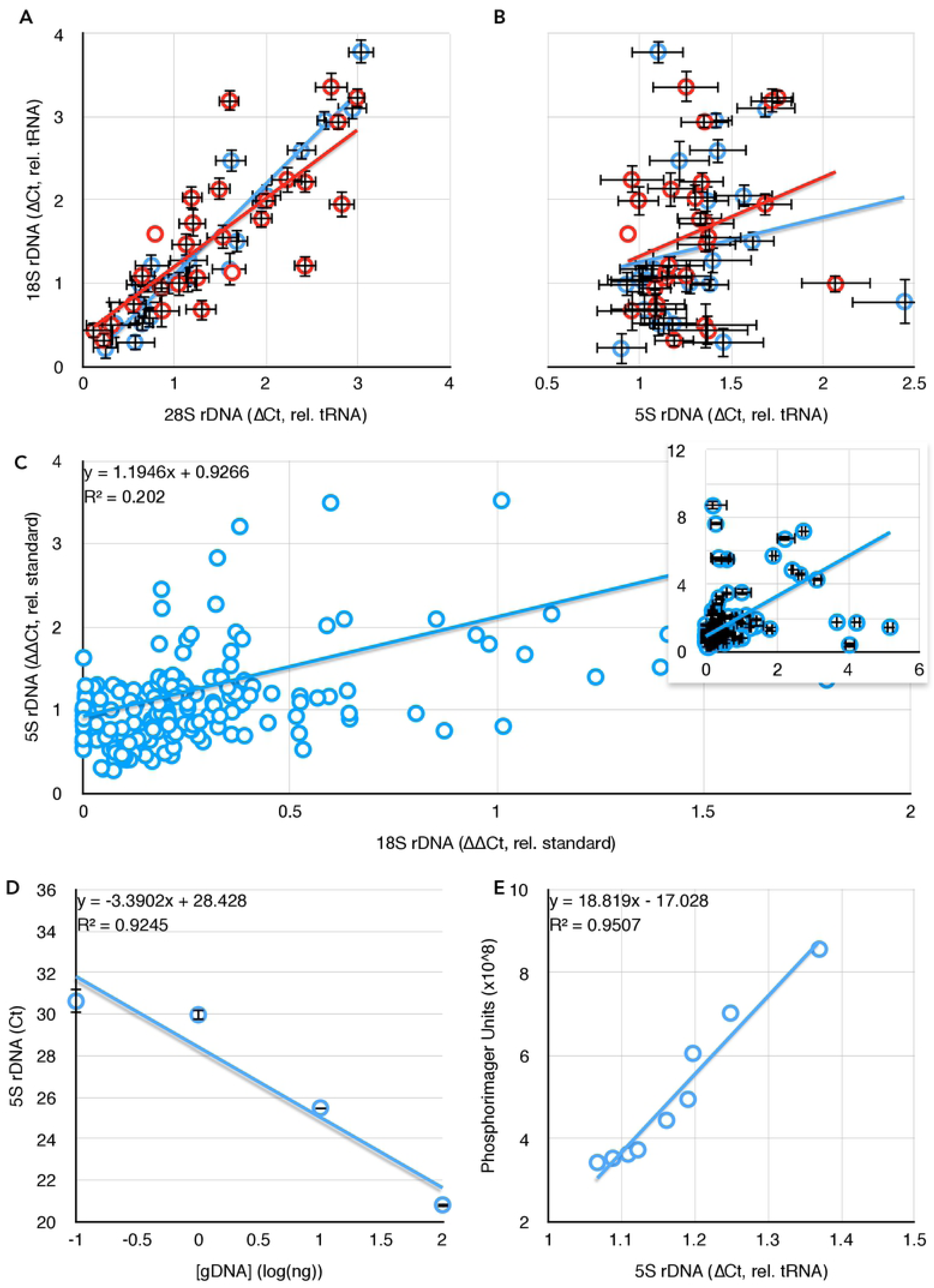
Comparisons of *18S rDNA* and *5S rDNA* in blood. **(A)** The correlation between *28S* and *18S* copy number is retained in blood samples taken from both women diagnosed with (red) or not diagnosed with breast cancer (blue). **(B)** The correlation between *5S* and *18S rDNA* copy numbers reported in (Gibbons et al. 2015) is not found in our dataset (no diagnosis, blue, R^2^ = 0.03; positive diagnosis, red, R^2^ = 0.08). (A) and (B) share an ordinate, error bars are S.E.M., and data are copy numbers relative to tRNA^Met^. **(C)** Extremely weak correlation between *18S rDNA* copy number and *5S* copy number in whole blood taken from people with no indication of any cancer diagnosis. Data are presented without error bars and with censoring of the highest values for clarity, but all data are present in the inset graph. **(D)** Template dose response for the *5S rDNA* primers, as in Figure 1B. **(E)** Comparison of *5S rDNA* copy number determined by qPCR and Southern blot quantification.

It is formally possible that the lack of *45S*-*5S* correlation in our samples is itself a pre-breast cancer indicator, and reflects an instability and loss of *5S* sequences in women at risk for breast cancer development. However, our results and those of Gibbons and colleagues (Gibbons et al. 2015) were derived from two different methodologies. Ours was *rDNA* copy number ascertainment from qPCR (relative to tRNA gene copy number), while the correlation reported by Gibbons and colleagues was derived from bioinformatic analysis of high throughput DNA sequencing, the real-time PCR validation data for which were not published. Therefore, the difference between *45S*-*5S* correlation in healthy and breast cancer patients could be a *bona fide* biomarker, or it could be an artifact of either method of determination.

### Quantification of *rDNA* Copy Number in Normal Human Blood

To address the correlation between *45S* and *5S rDNA* copy numbers in individuals not selected to have a history of breast cancer, we obtained 201 blood samples from the Arizona Health Sciences Center Biorepository at the Univeristy of Arizona (Table III). Too few healthy people donated blood for our needs, so we instead obtained blood from people of a broad range of age and both sexes whose blood was drawn as part of diagnosis of non-cancer afflictions. The blood was prepared as before, and *45S* and *5S* were determined relative to *tRNA*^*Met*^. In those samples, we observed the same lack of correlation between *45S* and *5S* copy number (Figure 6C). This adequately refutes our hypothesis that *rDNA* copy number differences could prefigure breast cancer development late in life, and further refutes the possibility that breast cancer development disrupts a natural coupling of *45S* and *5S* copy number. However, it does not identify the origin of the disparity between the data of Gibbons and colleagues and our own: it could be that informatic calculation of *rDNA* copy number is error-prone, or it could be that qPCR determination is insensitive, particularly with respect to the *5S*. We disfavor the latter possibility as we see no evidence of such when quantifying *5S* copy number over a range of genomic DNA concentrations (Figure 6D).

**Table III.**
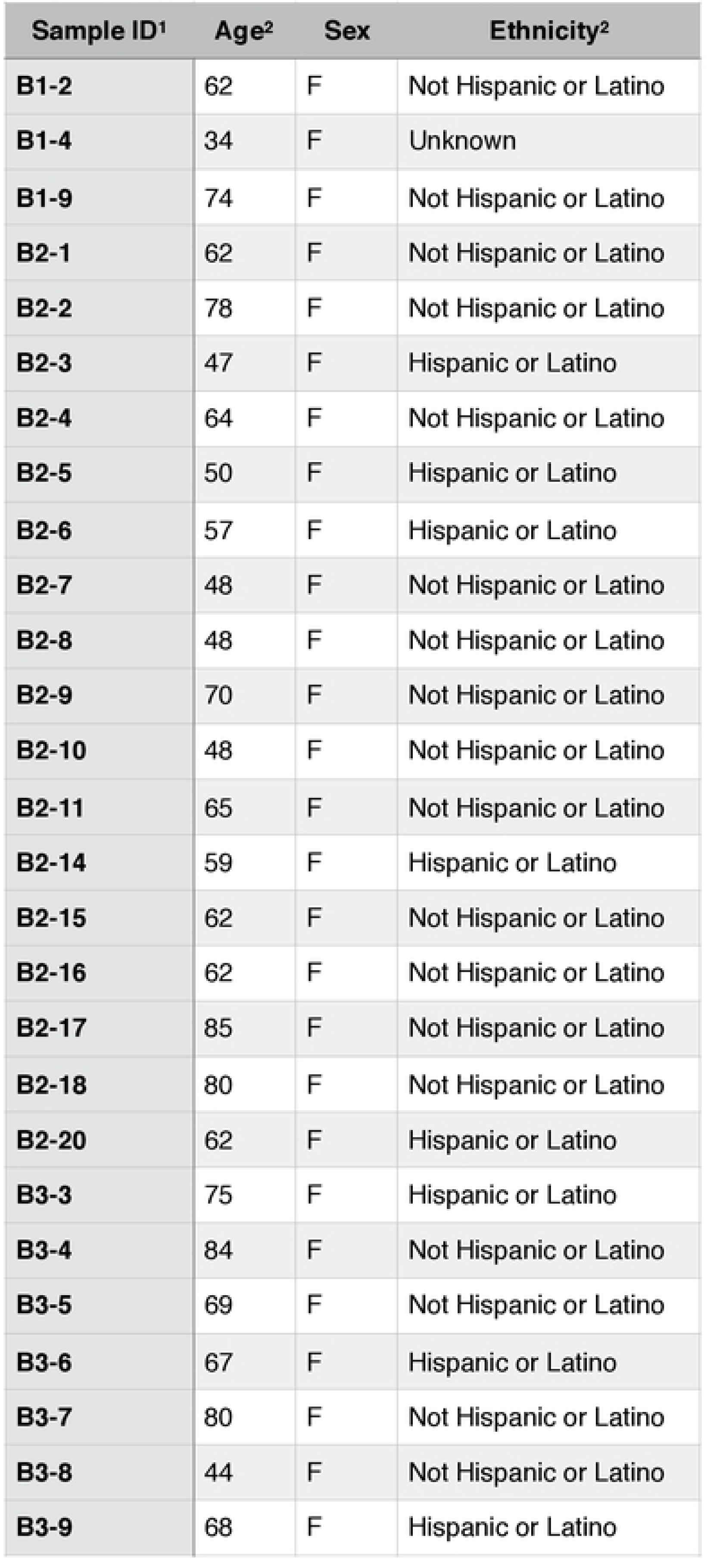
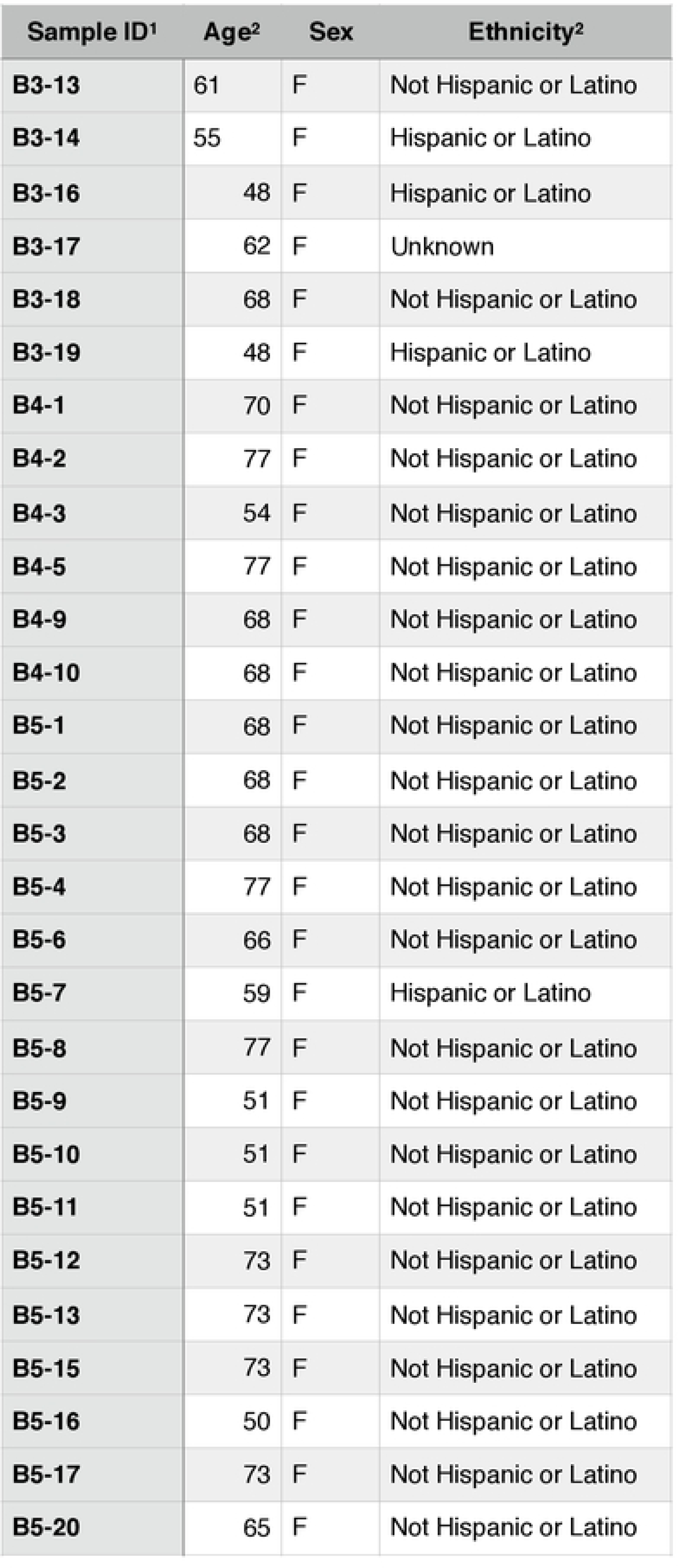
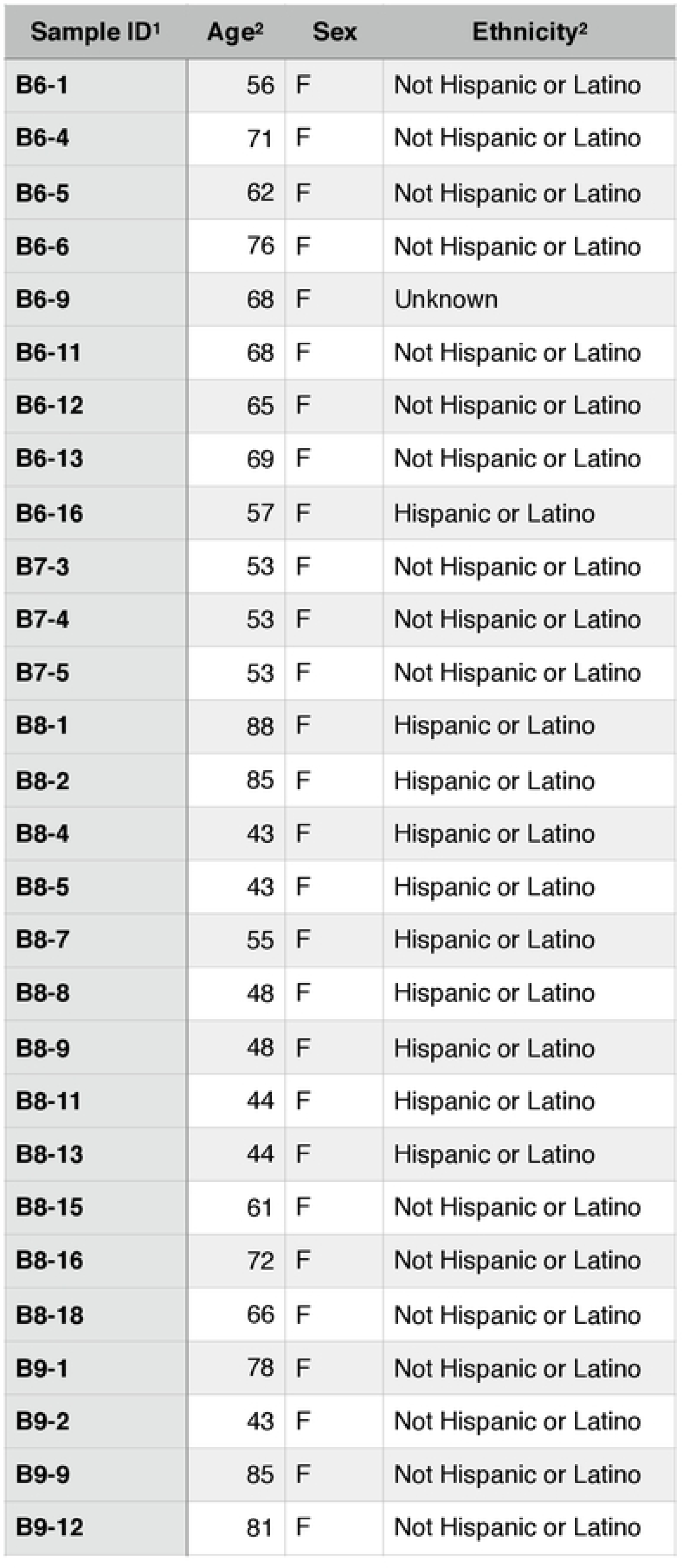
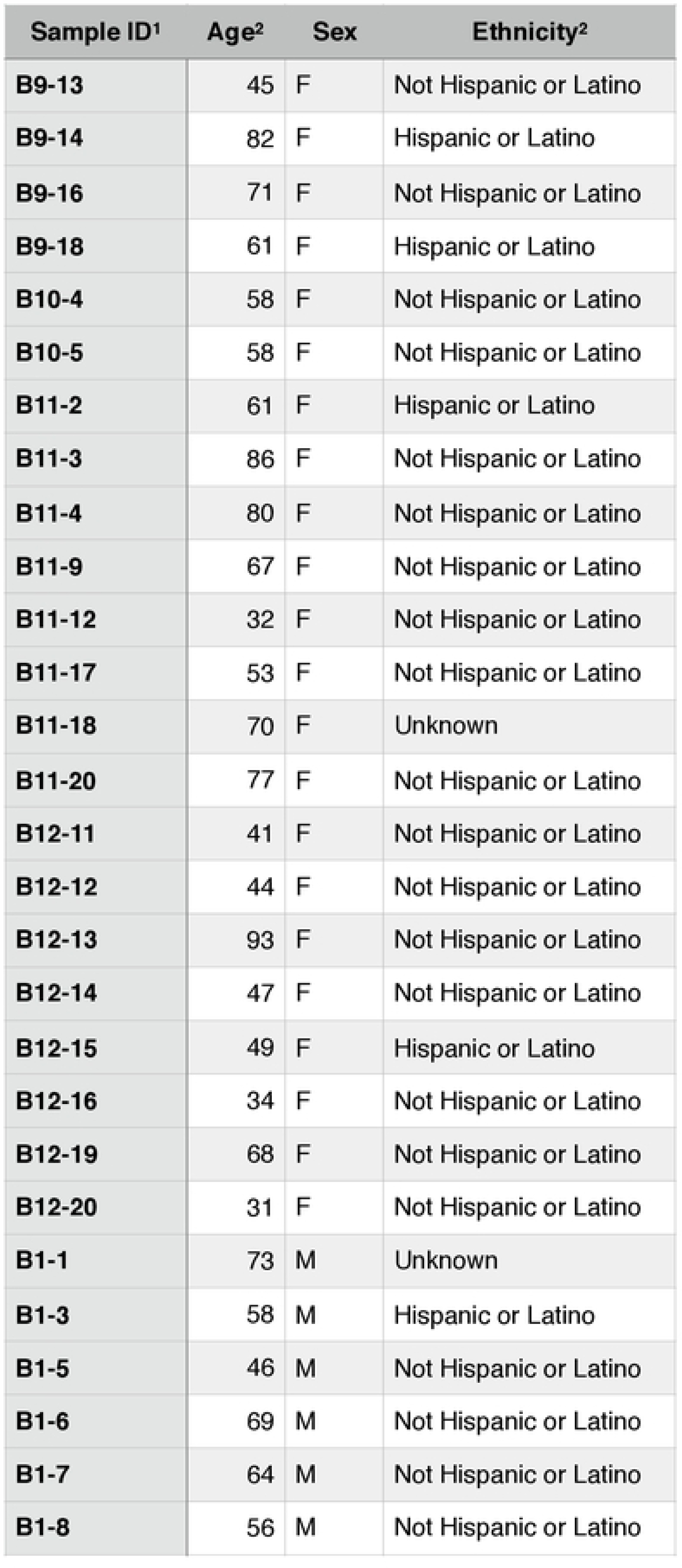
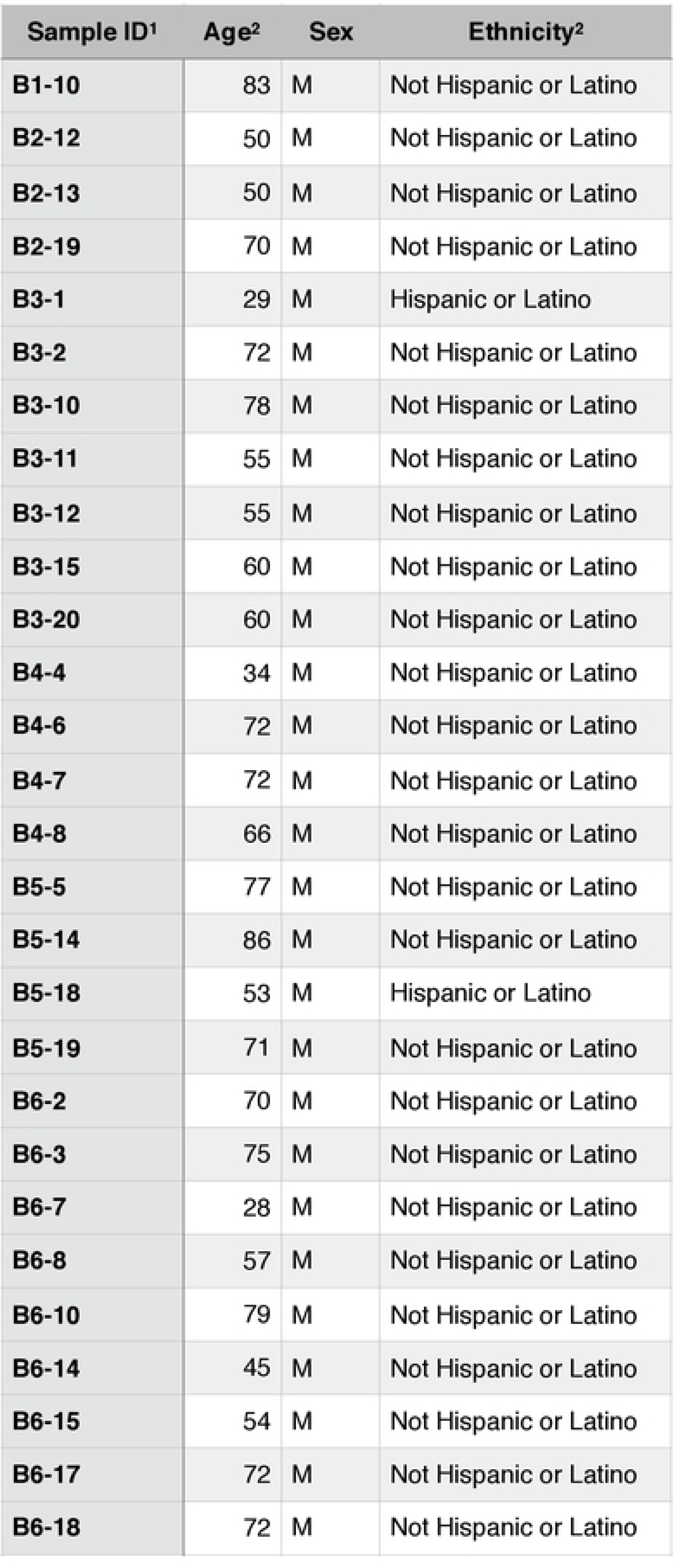
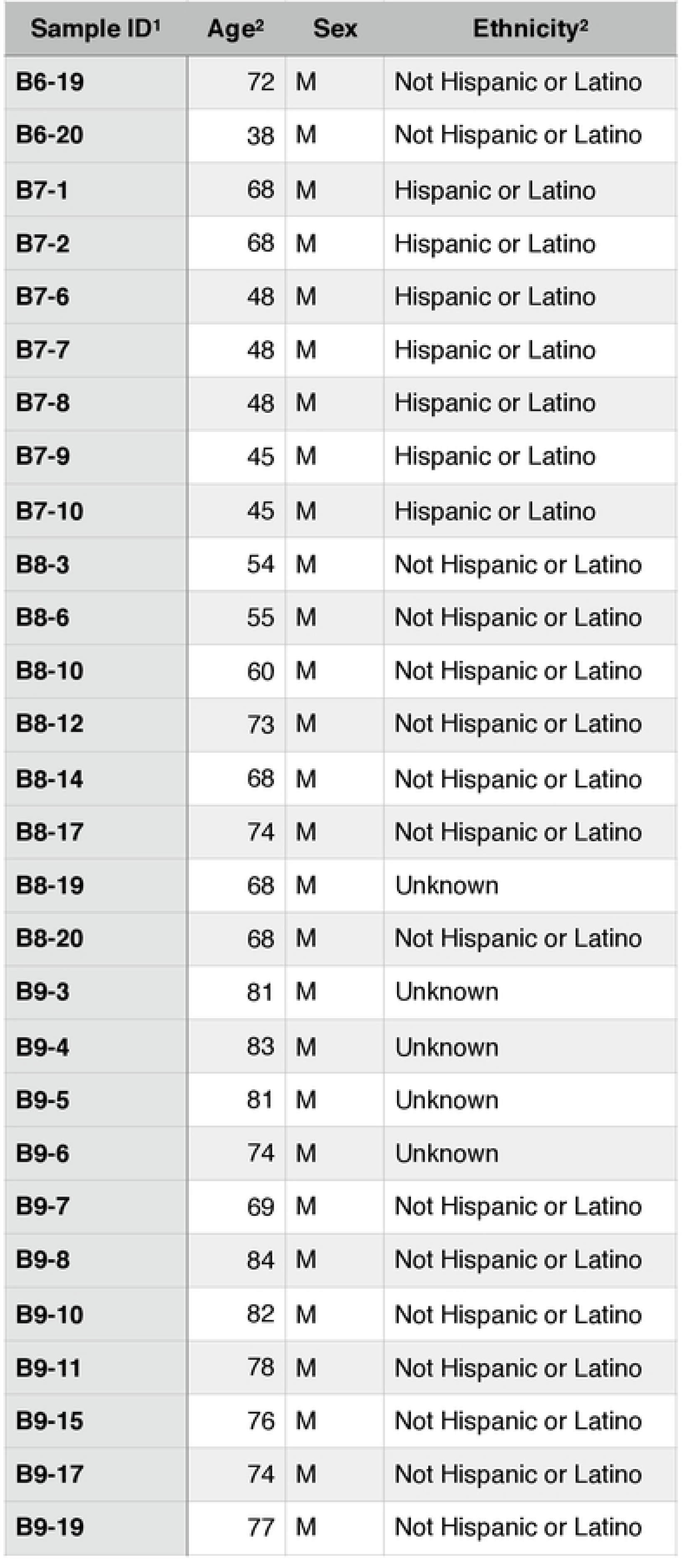
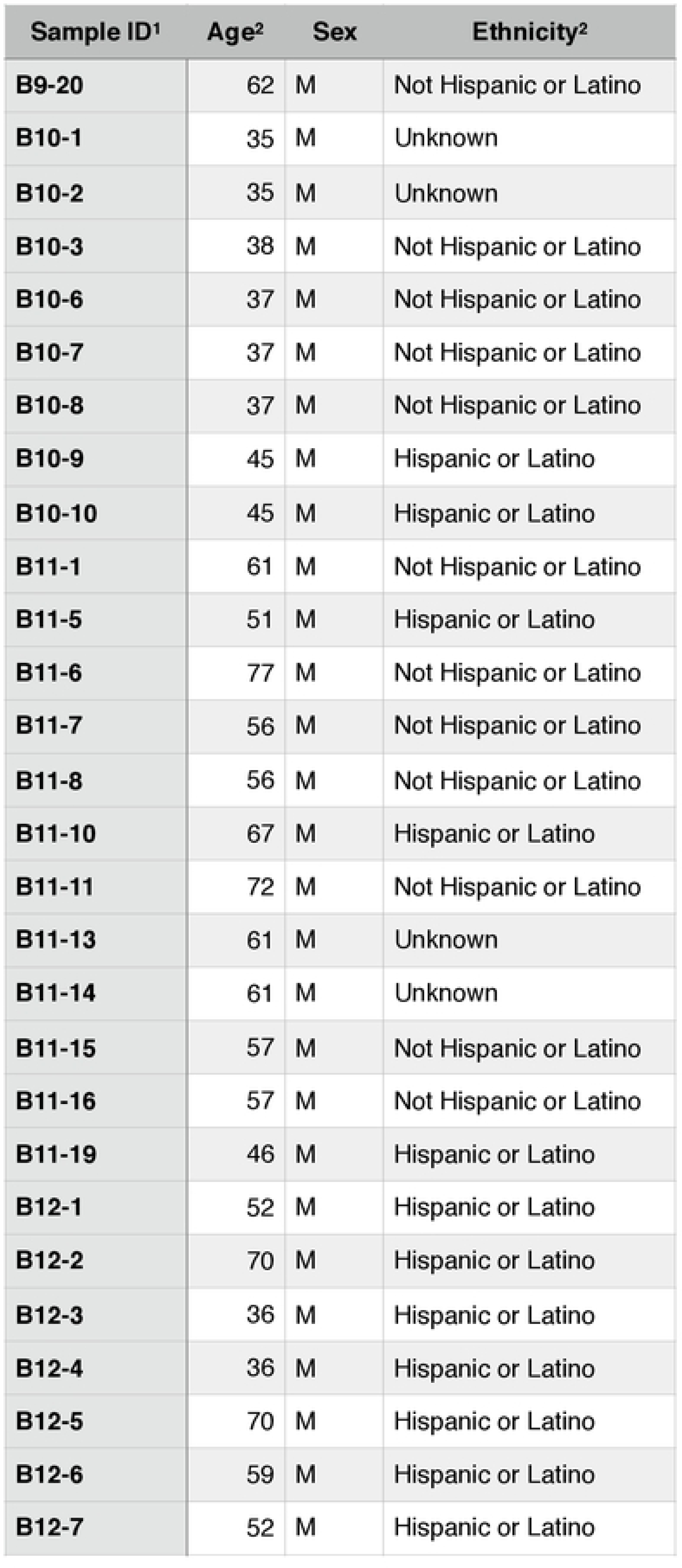
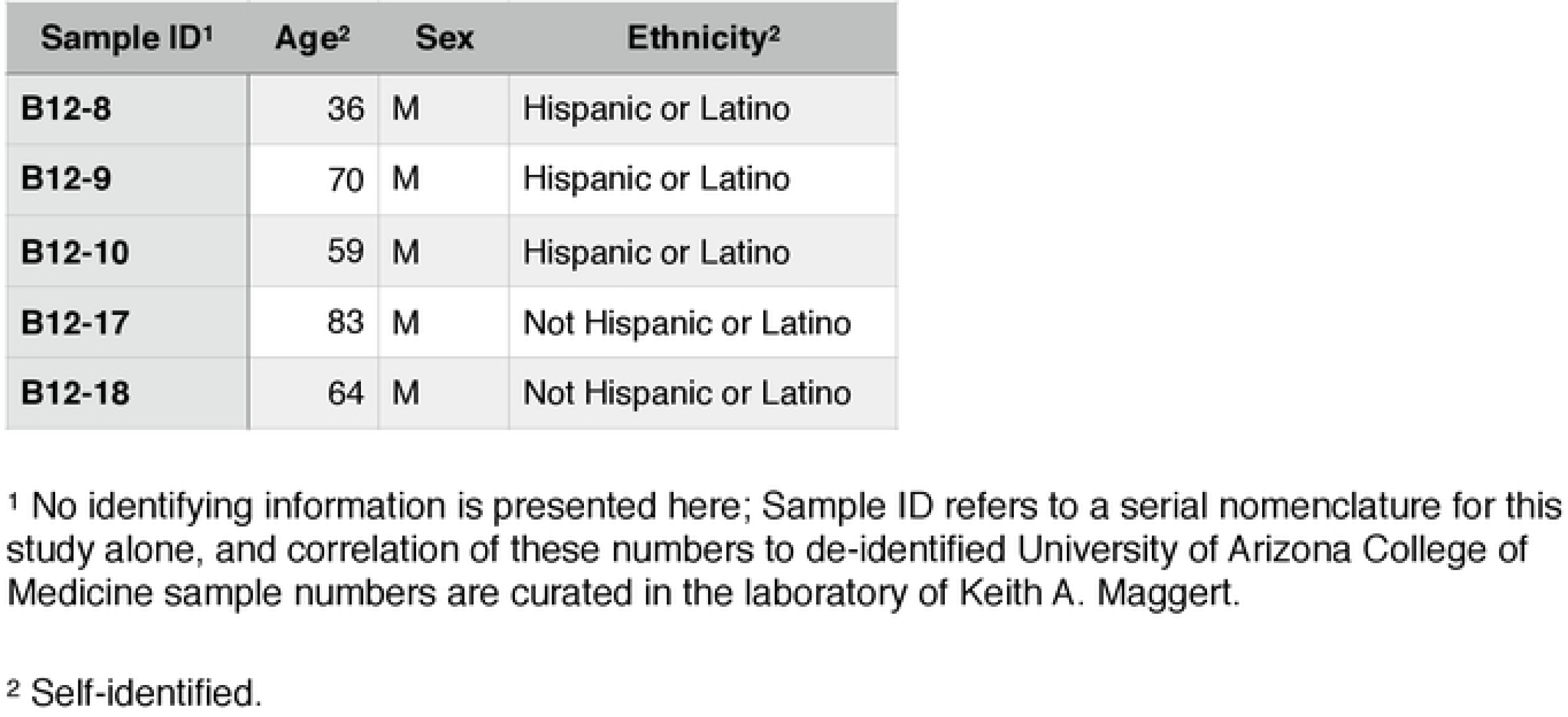
Demographics of blood samples

| **Sample ID<sup>1</sup>** | **Age<sup>2</sup>** | **Sex** | **Ethnicity<sup>2</sup>** |
| --- | --- | --- | --- |
| **B1-2** | 62 | F | Not Hispanic or Latino |
| **B1-4** | 34 | F | Unknown |
| **B1-9** | 74 | F | Not Hispanic or Latino |
| **B2-1** | 62 | F | Not Hispanic or Latino |
| **B2-2** | 78 | F | Not Hispanic or Latino |
| **B2-3** | 47 | F | Hispanic or Latino |
| **B2-4** | 64 | F | Not Hispanic or Latino |
| **B2-5** | 50 | F | Hispanic or Latino |
| **B2-6** | 57 | F | Hispanic or Latino |
| **B2-7** | 48 | F | Not Hispanic or Latino |
| **B2-8** | 48 | F | Not Hispanic or Latino |
| **B2-9** | 70 | F | Not Hispanic or Latino |
| **B2-10** | 48 | F | Not Hispanic or Latino |
| **B2-11** | 65 | F | Not Hispanic or Latino |
| **B2-14** | 59 | F | Hispanic or Latino |
| **B2-15** | 62 | F | Not Hispanic or Latino |
| **B2-16** | 62 | F | Not Hispanic or Latino |
| **B2-17** | 85 | F | Not Hispanic or Latino |
| **B2-18** | 80 | F | Not Hispanic or Latino |
| **B2-20** | 62 | F | Hispanic or Latino |
| **B3-3** | 75 | F | Hispanic or Latino |
| **B3-4** | 84 | F | Not Hispanic or Latino |
| **B3-5** | 69 | F | Not Hispanic or Latino |
| **B3-6** | 67 | F | Hispanic or Latino |
| **B3-7** | 80 | F | Not Hispanic or Latino |
| **B3-8** | 44 | F | Not Hispanic or Latino |
| **B3-9** | 68 | F | Hispanic or Latino |

| Sample ID<sup>1</sup> | Age<sup>2</sup> | Sex | Ethnicity<sup>2</sup> |
| --- | --- | --- | --- |
| B3-13 | 61 | F | Not Hispanic or Latino |
| B3-14 | 55 | F | Hispanic or Latino |
| B3-16 | 48 | F | Hispanic or Latino |
| B3-17 | 62 | F | Unknown |
| B3-18 | 68 | F | Not Hispanic or Latino |
| B3-19 | 48 | F | Hispanic or Latino |
| B4-1 | 70 | F | Not Hispanic or Latino |
| B4-2 | 77 | F | Not Hispanic or Latino |
| B4-3 | 54 | F | Not Hispanic or Latino |
| B4-5 | 77 | F | Not Hispanic or Latino |
| B4-9 | 68 | F | Not Hispanic or Latino |
| B4-10 | 68 | F | Not Hispanic or Latino |
| B5-1 | 68 | F | Not Hispanic or Latino |
| B5-2 | 68 | F | Not Hispanic or Latino |
| B5-3 | 68 | F | Not Hispanic or Latino |
| B5-4 | 77 | F | Not Hispanic or Latino |
| B5-6 | 66 | F | Not Hispanic or Latino |
| B5-7 | 59 | F | Hispanic or Latino |
| B5-8 | 77 | F | Not Hispanic or Latino |
| B5-9 | 51 | F | Not Hispanic or Latino |
| B5-10 | 51 | F | Not Hispanic or Latino |
| B5-11 | 51 | F | Not Hispanic or Latino |
| B5-12 | 73 | F | Not Hispanic or Latino |
| B5-13 | 73 | F | Not Hispanic or Latino |
| B5-15 | 73 | F | Not Hispanic or Latino |
| B5-16 | 50 | F | Not Hispanic or Latino |
| B5-17 | 73 | F | Not Hispanic or Latino |
| B5-20 | 65 | F | Not Hispanic or Latino |
| B6-1 | 56 | F | Not Hispanic or Latino |
| B6-4 | 71 | F | Not Hispanic or Latino |
| B6-5 | 62 | F | Not Hispanic or Latino |
| B6-6 | 76 | F | Not Hispanic or Latino |
| B6-9 | 68 | F | Unknown |
| B6-11 | 68 | F | Not Hispanic or Latino |
| B6-12 | 65 | F | Not Hispanic or Latino |
| B6-13 | 69 | F | Not Hispanic or Latino |
| B6-16 | 57 | F | Hispanic or Latino |
| B7-3 | 53 | F | Not Hispanic or Latino |
| B7-4 | 53 | F | Not Hispanic or Latino |
| B7-5 | 53 | F | Not Hispanic or Latino |
| B8-1 | 88 | F | Hispanic or Latino |
| B8-2 | 85 | F | Hispanic or Latino |
| B8-4 | 43 | F | Hispanic or Latino |
| B8-5 | 43 | F | Hispanic or Latino |
| B8-7 | 55 | F | Hispanic or Latino |
| B8-8 | 48 | F | Hispanic or Latino |
| B8-9 | 48 | F | Hispanic or Latino |
| B8-11 | 44 | F | Hispanic or Latino |
| B8-13 | 44 | F | Hispanic or Latino |
| B8-15 | 61 | F | Not Hispanic or Latino |
| B8-16 | 72 | F | Not Hispanic or Latino |
| B8-18 | 66 | F | Not Hispanic or Latino |
| B9-1 | 78 | F | Not Hispanic or Latino |
| B9-2 | 43 | F | Not Hispanic or Latino |
| B9-9 | 85 | F | Not Hispanic or Latino |
| B9-12 | 81 | F | Not Hispanic or Latino |
| B9-13 | 45 | F | Not Hispanic or Latino |
| B9-14 | 82 | F | Hispanic or Latino |
| B9-16 | 71 | F | Not Hispanic or Latino |
| B9-18 | 61 | F | Hispanic or Latino |
| B10-4 | 58 | F | Not Hispanic or Latino |
| B10-5 | 58 | F | Not Hispanic or Latino |
| B11-2 | 61 | F | Hispanic or Latino |
| B11-3 | 86 | F | Not Hispanic or Latino |
| B11-4 | 80 | F | Not Hispanic or Latino |
| B11-9 | 67 | F | Not Hispanic or Latino |
| B11-12 | 32 | F | Not Hispanic or Latino |
| B11-17 | 53 | F | Not Hispanic or Latino |
| B11-18 | 70 | F | Unknown |
| B11-20 | 77 | F | Not Hispanic or Latino |
| B12-11 | 41 | F | Not Hispanic or Latino |
| B12-12 | 44 | F | Not Hispanic or Latino |
| B12-13 | 93 | F | Not Hispanic or Latino |
| B12-14 | 47 | F | Not Hispanic or Latino |
| B12-15 | 49 | F | Hispanic or Latino |
| B12-16 | 34 | F | Not Hispanic or Latino |
| B12-19 | 68 | F | Not Hispanic or Latino |
| B12-20 | 31 | F | Not Hispanic or Latino |
| B1-1 | 73 | M | Unknown |
| B1-3 | 58 | M | Hispanic or Latino |
| B1-5 | 46 | M | Not Hispanic or Latino |
| B1-6 | 69 | M | Not Hispanic or Latino |
| B1-7 | 64 | M | Not Hispanic or Latino |
| B1-8 | 56 | M | Not Hispanic or Latino |
| **B1-10** | 83 | M | Not Hispanic or Latino |
| **B2-12** | 50 | M | Not Hispanic or Latino |
| **B2-13** | 50 | M | Not Hispanic or Latino |
| **B2-19** | 70 | M | Not Hispanic or Latino |
| **B3-1** | 29 | M | Hispanic or Latino |
| **B3-2** | 72 | M | Not Hispanic or Latino |
| **B3-10** | 78 | M | Not Hispanic or Latino |
| **B3-11** | 55 | M | Not Hispanic or Latino |
| **B3-12** | 55 | M | Not Hispanic or Latino |
| **B3-15** | 60 | M | Not Hispanic or Latino |
| **B3-20** | 60 | M | Not Hispanic or Latino |
| **B4-4** | 34 | M | Not Hispanic or Latino |
| **B4-6** | 72 | M | Not Hispanic or Latino |
| **B4-7** | 72 | M | Not Hispanic or Latino |
| **B4-8** | 66 | M | Not Hispanic or Latino |
| **B5-5** | 77 | M | Not Hispanic or Latino |
| **B5-14** | 86 | M | Not Hispanic or Latino |
| **B5-18** | 53 | M | Hispanic or Latino |
| **B5-19** | 71 | M | Not Hispanic or Latino |
| **B6-2** | 70 | M | Not Hispanic or Latino |
| **B6-3** | 75 | M | Not Hispanic or Latino |
| **B6-7** | 28 | M | Not Hispanic or Latino |
| **B6-8** | 57 | M | Not Hispanic or Latino |
| **B6-10** | 79 | M | Not Hispanic or Latino |
| **B6-14** | 45 | M | Not Hispanic or Latino |
| **B6-15** | 54 | M | Not Hispanic or Latino |
| **B6-17** | 72 | M | Not Hispanic or Latino |
| **B6-18** | 72 | M | Not Hispanic or Latino |

| Sample ID<sup>1</sup> | Age<sup>2</sup> | Sex | Ethnicity<sup>2</sup> |
| --- | --- | --- | --- |
| B6-19 | 72 | M | Not Hispanic or Latino |
| B6-20 | 38 | M | Not Hispanic or Latino |
| B7-1 | 68 | M | Hispanic or Latino |
| B7-2 | 68 | M | Hispanic or Latino |
| B7-6 | 48 | M | Hispanic or Latino |
| B7-7 | 48 | M | Hispanic or Latino |
| B7-8 | 48 | M | Hispanic or Latino |
| B7-9 | 45 | M | Hispanic or Latino |
| B7-10 | 45 | M | Hispanic or Latino |
| B8-3 | 54 | M | Not Hispanic or Latino |
| B8-6 | 55 | M | Not Hispanic or Latino |
| B8-10 | 60 | M | Not Hispanic or Latino |
| B8-12 | 73 | M | Not Hispanic or Latino |
| B8-14 | 68 | M | Not Hispanic or Latino |
| B8-17 | 74 | M | Not Hispanic or Latino |
| B8-19 | 68 | M | Unknown |
| B8-20 | 68 | M | Not Hispanic or Latino |
| B9-3 | 81 | M | Unknown |
| B9-4 | 83 | M | Unknown |
| B9-5 | 81 | M | Unknown |
| B9-6 | 74 | M | Unknown |
| B9-7 | 69 | M | Not Hispanic or Latino |
| B9-8 | 84 | M | Not Hispanic or Latino |
| B9-10 | 82 | M | Not Hispanic or Latino |
| B9-11 | 78 | M | Not Hispanic or Latino |
| B9-15 | 76 | M | Not Hispanic or Latino |
| B9-17 | 74 | M | Not Hispanic or Latino |
| B9-19 | 77 | M | Not Hispanic or Latino |
| B9-20 | 62 | M | Not Hispanic or Latino |
| B10-1 | 35 | M | Unknown |
| B10-2 | 35 | M | Unknown |
| B10-3 | 38 | M | Not Hispanic or Latino |
| B10-6 | 37 | M | Not Hispanic or Latino |
| B10-7 | 37 | M | Not Hispanic or Latino |
| B10-8 | 37 | M | Not Hispanic or Latino |
| B10-9 | 45 | M | Hispanic or Latino |
| B10-10 | 45 | M | Hispanic or Latino |
| B11-1 | 61 | M | Not Hispanic or Latino |
| B11-5 | 51 | M | Hispanic or Latino |
| B11-6 | 77 | M | Not Hispanic or Latino |
| B11-7 | 56 | M | Not Hispanic or Latino |
| B11-8 | 56 | M | Not Hispanic or Latino |
| B11-10 | 67 | M | Hispanic or Latino |
| B11-11 | 72 | M | Not Hispanic or Latino |
| B11-13 | 61 | M | Unknown |
| B11-14 | 61 | M | Unknown |
| B11-15 | 57 | M | Not Hispanic or Latino |
| B11-16 | 57 | M | Not Hispanic or Latino |
| B11-19 | 46 | M | Hispanic or Latino |
| B12-1 | 52 | M | Hispanic or Latino |
| B12-2 | 70 | M | Hispanic or Latino |
| B12-3 | 36 | M | Hispanic or Latino |
| B12-4 | 36 | M | Hispanic or Latino |
| B12-5 | 70 | M | Hispanic or Latino |
| B12-6 | 59 | M | Hispanic or Latino |
| B12-7 | 52 | M | Hispanic or Latino |
| **B12-8** | 36 | M | Hispanic or Latino |
| **B12-9** | 70 | M | Hispanic or Latino |
| **B12-10** | 59 | M | Hispanic or Latino |
| **B12-17** | 83 | M | Not Hispanic or Latino |
| **B12-18** | 64 | M | Not Hispanic or Latino |

<sup>1</sup> No identifying information is presented here; Sample ID refers to a serial nomenclature for this study alone, and correlation of these numbers to de-identified University of Arizona College of Medicine sample numbers are curated in the laboratory of Keith A. Maggert.
<sup>2</sup> Self-identified.

To resolve this conflict, we selected blood samples with large and small *5S rDNA* counts and subjected them to Southern dot-blot analysis. We observed a very strong correlation between Southern-based copy number determination and the results from qPCR across a broad range (Figure 6E). This is in good agreement with similar experiments in *Drosophila*, where very small differences in *18S rDNA* copy number by qPCR could be robustly quantified and then independently validated by genetic means (Paredes and Maggert 2009a). Thus, our findings question the value of *rDNA* copy number determination using DNA deposited in high-throughput sequencing databases, consistent with those reported *rDNA* counts being so at-odds with other methodologies (Long and Dawid 1980). We cannot speculate whether any potential error that may exist in databases is a result of biases in sequencing, quality-control, the extraction and analysis of copy number data, or some other step in the informatics “pipeline.” It seems *prima facie* erroneous that some samples had fewer than 10 copies of either *45S* or *5S* rDNAs (Wang and Lemos 2017), which is incompatible with life in any known eukaryote (Long and Dawid 1980; Prokopowich et al. 2003). Instead, our data support our assertion that the existing databases are not reliable sources of *rDNA* copy number data, and it is likely that if *rDNA* copy number proves to be a risk factor or a diagnostic for human disease, then a simple and reliable protocol, such as ours, will be necessary to validate data in databases or from patients.

### *rDNA* Copy Numbers Do Not Correlate with Age, Sex, or Ethnicity

Our collection of blood from breast cancer patients, and from patients with non-cancer diagnoses, allowed us to ascertain deviations in average *rDNA* copy number as a consequence of age, sex, and ethnicity. These data could prove useful in evaluating hypotheses linking *rDNA* copy number to age (Malinovskaya et al. 2018; Wang and Lemos 2019), disease status, or involvement in lifestyle choices that lead to greater disease risk (*e.g.*, alcohol consumption, cigarette smoking, overeating (Holland et al. 2016)). We re-analyzed the *rDNA* copy number from the non-cancer-diagnosis individuals from the Tucson area and this time looked for correlations between average *rDNA* copy number and sex and ethnicity as categorical conditions (Figure 7A-B). No condition grouped *rDNA* copy number significantly away from the others. Thus, we conclude that in our samples, there is no evidence to support the hypothesis that *rDNA* copy number differs in the blood of different sexes or ethnicities.

**Figure 7.**
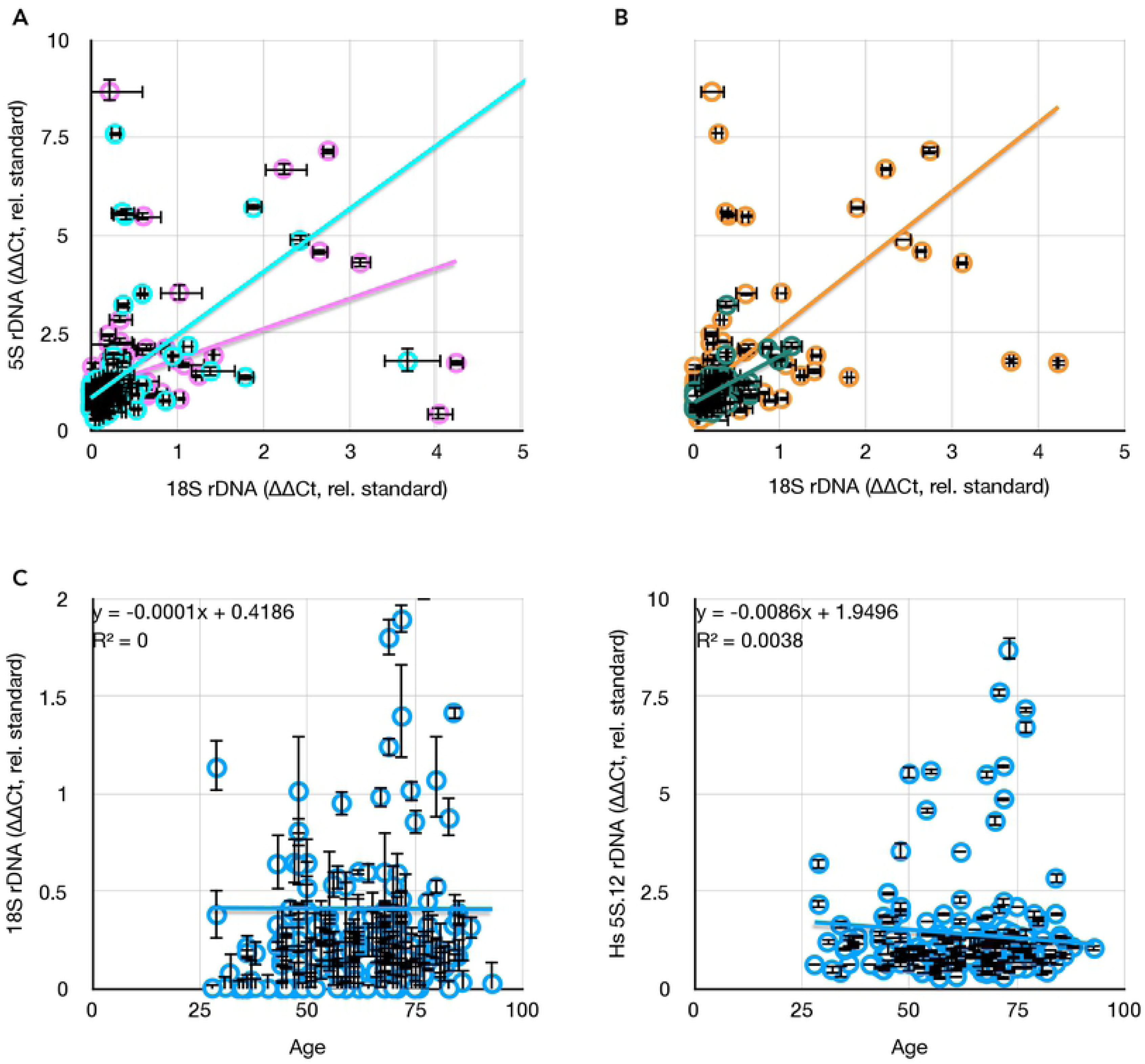
Comparisons between *rDNA* copy numbers and population variables. **(A)** Consistent lack of correlation between different *rDNA* cluster copy numbers (*18S* and *5S*) in males (blue, y = 1.61x + 0.83, R^2^ = 0.24) and females (pink, y = 0.78x + 1.06, R^2^ = 0.19). **(B)** Consistent weak correlation between different *rDNA* cluster copy numbers (*18S* and *5S*) in hispanic (green, y = 1.20x + 0.72, R^2^ = 0.37) and non-hispanic white (orange, y = 1.75x + 0.87, R^2^ = 0.30) individuals. **(C)** Consistent lack of correlation between *18S* and *5S rDNA* copy numbers as a function of age of blood donor. All values in this graph are represented as *rDNA* copy number relative to tRNA^Met^, and normalized to the lab standard.

Aging has been linked to *rDNA* loss in yeast, and the application of that phenomenon to humans has been proposed and, in some cases, supported (Malinovskaya et al. 2018). In contrast to those studies, we could detect no correlation between individual *rDNA* copy number or C.V. with age at time of blood collection (Figure 7C). This lack of apparent correlation is no-doubt affected by the widely-variant copy number at birth of these patients, and it is still possible that individuals will exhibit *rDNA* loss as a function of aging. At present, sufficiently broad sampling of blood over the lifetimes of individuals is not available, so while the issue of individual *rDNA* copy number loss is still outstanding, it can be said that “snapshot” *rDNA* copy number is not predictive of biological age. This alone makes *rDNA* copy number a poor metric for age or sex, and extreme caution must be employed before considering such metrics in disease risk, disease onset, or forensic identification.

### Concluding Remarks

We have presented a method for quickly, inexpensively, sensitively, accurately, and reproducibly determining the copy number of repeated DNAs, in particular the ribosomal RNA genes (*45S* and *5S rDNA*s), from patient samples. This approach was pioneered in the model system *Drosophila*, but it’s application here has allowed us to directly test multiple hypotheses concerning repeat DNA stability in both healthy individuals and breast tumors.

We find, first, that the *rDNA* show signs of general instability, consistent with previous work showing derepression of heterochromatin-mediated stability in cancer. This finding is supported by our demonstration that the satellite repeats vary in time with (although not in the same direction as) increased *rDNA* variation. Second, we find that the reported covariation between *45S* and *5S* gene copy numbers is not evident using techniques other than high-throughput sequencing. Our data demonstrating the sensitivity, responsiveness, and confirmation by Southern blot analyses suggests that great caution must be exercised when deriving repeat copy numbers from curated sequencing data. Third, our findings refute hypotheses suggesting that increases or decreases in *rDNA* copy number are adaptive for disease. Rather than being selected for by increased demands for protein synthesis, or selected against by trimming genomes of superfluous DNA, we conclude that hyper-variability is a general outcome of disease onset. It is likely that the observed preferential losses in cultured cancer cell lines are an artifact of growth *in vitro*, and that losses may merely be more stable than gains in culture. Fourth, we have demonstrated a facile, rapid, inexpensive, precise, and accurate real time PCR based strategy for *rDNA* copy number quantification using very small amounts of fresh or fixed tissue.

## Acknowledgements

GFP-BLM cell lines were obtained from Dr. Mary Yagle and Dr. Nathan Ellis. The University of Arizona Cancer Center provided core and facilities support, funded through National Institutes of Health Support Grant P30CA023074. Blood DNA (Figures 6A-B, E) or blood samples (Figures 6C and 7) were obtained from the AHSC Biorepository at The University of Arizona College of Medicine. Ovary samples (Figure 2) were obtained from the Univeristy of Arizona Tissue Acquisition and Cellular/Molecular Analysis Shared Resource (TACMASR). TACMASR also performed the tissue sections of the breast tumor samples. Support also came from the Department of Cellular and Molecular Medicine, the Univeristy of Arizona College of Medicine, the Arizona State Museum, and Transformative Research Award (R01) GM123640, granted to Keith A. Maggert. Finally, Dr. Nicholas Ratterman’s excellent sociolinguistic parsimony and keen advice was invaluable, as were the delicate urgings of Dr. Diana Darnell.

## Materials and Methods

### DNA from Tumor and Tissue Samples

Five consecutive 1 *µ*m slices were cut from formalin-fixed-paraffin-embedded tissue blocks and DNA purification was obtained by using QIAamp DNA FFPE Tissue Kit (Qiagen). Basically, 1 ml of xylenes was added to each tube, vortexed for 1 minute, then centrifuged at 14000 rpm for 2 minutes and the supernatant removed through two ethanol washes. For each, ATL buffer and proteinase K were added, the samples ground with a pestle, and incubated for 1 hour at 56C° and 1 hour at 90C°. RNAse A was added to the samples and left at room temperature for 2 minutes, then the samples were transferred to columns and were washed with buffer AL, buffer AW1, and finally buffer AW2. Samples were eluted with 100 *µ*L buffer ATE and quantified using a Synergy H1 Microplate Reader (BioTek).

### DNA from Blood Cards

DNA purification was performed using GenSolve DNA Recovery Kit (Gentegra), the QIAshredder Kit (Qiagen), and the QIAamp Blood Mini Kit (Qiagen). Basically, half of each blood card was cut and placed in individual microcentrifuge tubes with Recovery Solution A and the blood allowed to resuspend overnight at room temperature. Tubes were then incubated, with rotation, for 1 hour at 56C°. Recovery Solution B was added along with the samples to QIAshredder columns and centrifuged at 13300 rpm for 2 minutes. The columns were discarded and ethanol added, the samples vortexed and briefly centrifuged, then transferred to QIAamp columns and centrifuged at 8000 rpm for 1 minute. Columns were washed with buffer AW1 then with buffer AW2. Finally, samples were eluted in two steps of buffer AE, in a total of 100 μL of DNA solution, then quantified using a Synergy H1 Microplate Reader (BioTek).

### DNA from Drosophila and Human Blood

Tissue was ground with a mini-pestle in a microcentrifuge tube in a solution containing 100 mM Tris pH 8.0, 50 mM ethylenediaminetetraacetic acid, 1% Sodium Dodecylsulfate, and 1 *µ*g Proteinase K. The slurry was digested for an hour at 65°C, then extracted through a series of phenol, phenol-chloroform, chloroform, and ether. The DNA was ethanol-precipitated and resuspended in TE (10 mM Tris pH 8.0, 1 mM ethylenediaminetetraacetic acid) containing 0.1 *µ*g RNAseA. DNA concentration was determined using a Synergy H1 Microplate Reader (BioTek), and DNA samples were stored at high concentration at −80°C until use.

### Real-Time PCR Reactions

Real-Time PCR reactions were done as described previously (Paredes and Maggert 2009a; Aldrich and Maggert 2014), including controls on an ABI Step-One or Stop-One-Plus using SYBR Green chemistry, full-length (2 hour) reaction cycle, and obligate post-hoc melt curve analysis. Reactions were adjusted to be 12 *µ*L total reaction volume. Reactions were performed in triplicate or more, and data were accepted as valid if the standard error of the mean of the replicates was less than 0.1. Primers for non-*rDNA* targets are: *Bloom Helicase* (GGCTGCTGTTCCTCAAAATAATCTACAG and ATTATTAAGTGTTCTGGCTGAGTGACG), *Snail2* (CCCGTATCTCTATGAGAGTTACTCC and GTATGCTCCTGAGCTGAGGATCTC), *RNMT2* (CTTTGATGGCAGCATACAGTGTTCTGG and CCTGTGAATTTCTTCTGCAGTTTCAAGC), *eGFP* (GAGGGTGAAGGTGATGCAACATACGG and GCCATGGAACAGGTAGCTTCCCAG).

### Numerical Analysis

Analyses were performed on an Apple MacBook Pro using Numbers version 6.0 (build 6194). Descriptive and frequentist statistical analyses (*e.g.*, regression, slope/intercept) were from embedded functions.

For simple comparisons between copy numbers of two genes in the same sample (*e.g.*, Figure 1B, 1C), Crossing thresholds (C_t_s) could be directly compared. C_t_ is calculated by determining the PCR cycle at which the signal first crosses the average plus 10 standard deviations of all preceding cycles. Errors are generally presented as Standard Errors of the Mean, derived from the pooled errors of the copy number of the gene in question and the tRNA normalizer, run in triplicate or quadruplicate. Standard error of the mean is justified as these data are from technical (assay) replicates of DNA extracted from single individuals. Errors were pooled using the standard summation of errors 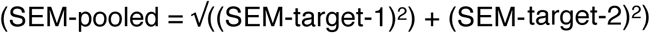.

For copy number determination between different samples (*e.g.*, Figure 1D, 5A-C) crossing thresholds were converted to relative amounts by first subtracting tRNA C_t_ from the target gene C_t_, then using that as an exponent (2^(Ct-target – Ct-tRNA)^; this is commonly referred to as the “ΔC_t_” method. Where appropriate (*e.g.*, Figures 6C, 7), the “ΔΔC_t_” method was used, which calculates ΔC_t_s for the sample and for a standard (in this case a human DNA sample that is a study-wide standard in the laboratory); these data indicate the proportion of *rDNA*-to-tRNA in the sample relative to the *rDNA*-to-tRNA of the standard, allowing us to compare values between qPCR reactions.

Correlations were considered weak if R^2^ was below 0.65, and non-existent if below 0.20. *A priori* criteria for accepting Type-I errors (alpha) was set at 0.01 at the beginning of the study.

### Human Data

Breast cancer tumor and adjacent samples from 29 individuals were obtained from Dr. L. LeBeau. It was determined by the University of Arizona Institutional Review Board that the use of the tissues did not require board oversight for this study (protocol #15-0477-0333) as the work did not meet the definition of “human subjects” by U.S. Department of Health and Human Services which state that “human subject means a living individual about whom an investigator (whether professional or student) conducting research obtains data through intervention or interaction with the individual, or identifiable private information.” Samples were de-identified, and shared without personal information for the purpose of this study.

DNA samples from 51 individuals taking part in a trial to identify cancer risk alleles were collected under protocol #12-0138 (to C. Laukaitis), which was approved by the University of Arizona Institutional Review Board. Samples were de-identified, and shared without personal information for the purpose of this study.

Blood samples from 200 individual patients obtained from the Arizona Health Sciences Center Biorepository (from D. Harris) were de-identified, and shared without personal information for the purpose of this study.

Samples were further de-identified by assigning new serial numbers to all samples; the key linking the original de-identified number from the source (*i.e.*, LeBeau, Laukaitis, Harris) to the tables in this study is safeguarded by our laboratory.

